# The Critical Role of the C-terminal Lobe of Calmodulin in Activating Eukaryotic Elongation Factor 2 Kinase

**DOI:** 10.1101/2025.05.13.653565

**Authors:** Kimberly J. Long, Luke S. Browning, Andrea Piserchio, Eta A. Isiorho, Mohamed I. Gadallah, Jomai Douangvilay, Elizabeth Y. Wang, Justin K. Kalugin, Jennifer S. Brodbelt, Ranajeet Ghose, Kevin N. Dalby

## Abstract

Eukaryotic elongation factor-2 kinase (eEF-2K), a member of the α−kinase family, modulates translational rates by phosphorylating eEF-2, a GTPase that facilitates the translocation of the nascent chain on the ribosome during the elongation phase of protein synthesis. eEF-2K is regulated by diverse cellular cues, many of which sensitize it to the Ca^2+^-effector protein calmodulin (CaM). CaM, which binds and allosterically activates eEF-2K in the presence of Ca^2+^, contains two structural “lobes,” each with a pair of Ca^2+^-binding EF-hands. Using kinetic analysis, we demonstrate that the isolated C-terminal lobe of CaM (CaM_C_) is sufficient to engage and fully activate eEF-2K in a Ca^2+^-dependent fashion. Genetically fusing CaM_C_ to the N-terminus of eEF-2K, upstream of its critical CaM-targeting motif (CTM) via a flexible 2-glycine linker, results in a chimeric species (C-LiNK) that is constitutively active independent of external CaM and Ca^2+^. A structure of the C-LiNK functional core reveals no significant deviation in the overall conformations of the interacting modules and orientations of key catalytic-site residues relative to the heterodimeric complex between full-length CaM and eEF-2K. These observations demonstrate that, in contrast to other CaM-regulated kinases, CaM_C_ alone is sufficient to activate eEF-2K fully. The proximity effect of CaM_C_ in the context of C-LiNK removes the requirement for external Ca^2+^, whose apparent role is to enhance the CaM-affinity of eEF-2K and drive kinase activation. The responsiveness of eEF-2K to regulatory stimuli in cells appears to be lost in C-LiNK, presumably due to its permanently “on” state.

## Introduction

In eukaryotic cells, the GTPase eukaryotic elongation factor 2 (eEF-2) promotes the GTP-dependent translocation of the nascent chain from the ribosomal amino-acyl site to the peptidyl site during translational elongation. The ability of eEF-2 to associate with the ribosome is regulated by phosphorylation at Thr-56, which is mediated by the atypical serine/threonine kinase eukaryotic elongation factor 2 kinase (eEF-2K) (1–5), a member of the α−kinase family (6, 7). The activity of eEF-2K is finely tuned to cellular energy levels, nutrient availability, and environmental stress, allowing it to dynamically regulate protein synthesis in response to specific cellular cues (8–10). It has been demonstrated that fundamental processes like memory formation and stress adaptation rely on changes in elongation rates, which play a crucial role in translational reprogramming (11, 12). This adaptive process prioritizes the translation of specific mRNAs by modulating the corresponding rates relative to global translation. Given its pivotal role in regulating protein synthesis, it is not surprising that dysregulated eEF-2K activity correlates with various diseases, including Alzheimer’s-related dementia (13), Parkinson’s disease (14), and tumorigenesis (15, 16).

eEF-2K is activated by the Ca^2+^-effector protein calmodulin (CaM) (2, 17) through a unique allosteric mechanism (18, 19) that sets it apart from other CaM-regulated kinases (20, 21). Following CaM binding, eEF-2K undergoes rapid autophosphorylation at an activating site (T348, Fig. 1) within its regulatory loop (R-loop, Fig. S1). The engagement of phosphorylated T348 (*p*T348) at a basic phosphate binding pocket (PBP) yields a conformational state capable of efficiently phosphorylating eEF-2. Recent crystal structures of the complex between CaM and a minimal CaM-activatable core of eEF-2K (eEF-2K_TR_, Fig. S1) phosphorylated at T348 (*p*eEF-2K_TR_) provide valuable structural insight into the unique activation mechanism of eEF-2K (18, 22, 23). The structures suggest intimate interactions between the Ca^2+^-loaded C-terminal lobe of CaM (CaM_C_) and eEF-2K; the N-terminal lobe of CaM (CaM_N_) does not appear to make any stable contacts within the complex. This mode of interaction is consistent with changes in the degree of protection seen for CaM_N_ and CaM_C_ upon formation of the CaM•*p*eEF-2K_TR_ complex through hydrogen/deuterium exchange mass spectrometry (HDX-MS) analyses (24).

**Fig. 1.**
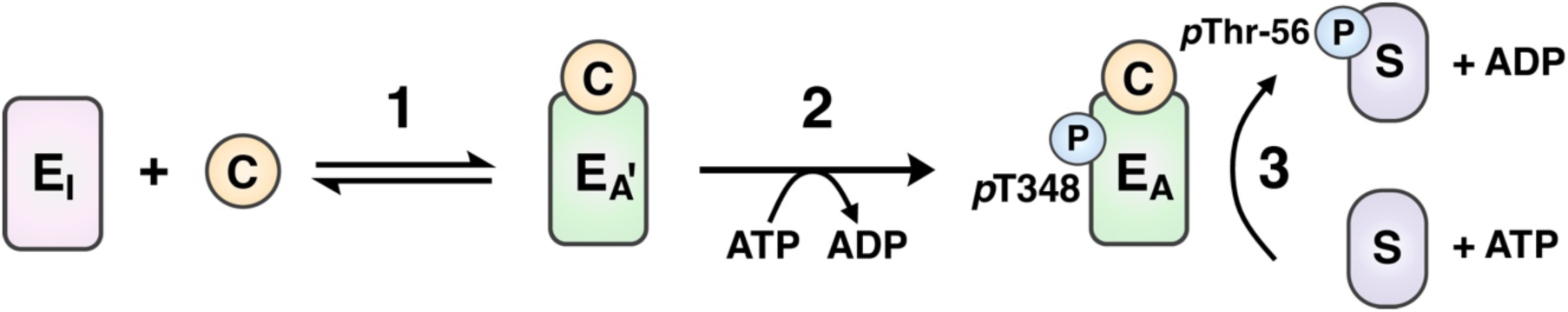
Schematic representation of the CaM-mediated activation of eEF-2K. CaMactivates eEF-2K through a two-step process. In the first step (1), CaM binds the eEF-2K that is in the inactive state (E_I_), leading to a state (E_A’_) that has high activity toward T348. In the second step (2), rapid autophosphorylation in T348 and its subsequent engagement in a phosphate binding leads to a fully activated state (E_A_) that can phosphorylate the substrate eEF-2 (S) on Thr-56 with high efficiency.

While structural data highlight the importance of CaM_C_ in the eEF-2K complex, a thorough biochemical assessment of the lobe-specific role of CaM in driving eEF-2K activation and activity is needed in the context of the intact enzyme *in vitro* and in cells. For example, the absence of a significant fraction of the dynamic regulatory loop (R-loop) in the eEF-2K_TR_ construct (Fig. S1) may result in the loss of specific regulatory interactions, potentially with CaM_N_. To investigate the contributions of the individual lobes of CaM in eEF-2K function, we assessed the ability of individual constructs encoding CaM_N_ or CaM_C_ to engage and activate wild-type eEF-2K in a Ca^2+^-sensitive fashion. We found that while the isolated CaM_N_ does not significantly stimulate eEF-2K activity, CaM_C_ displays a robust ability to bind and activate the enzyme in a Ca^2+^-dependent manner. By tethering CaM_C_ to the N-terminus of eEF-2K, we generated a constitutively active chimeric species insensitive to exogenous Ca^2+^. Additionally, this chimeric construct expressed in MCF10A^eEF-2K-/-^ cells displayed high activity even in the presence of typically suppressive post-translational modifications (PTMs). These findings support a model wherein CaM, specifically CaM_C_, acts as an essential allosteric activator of eEF-2K, with regulatory signals, including Ca^2+^ and diverse PTMs, dynamically modulating its interaction with eEF-2K to regulate its activity.

## Results

### CaM_C_ drives eEF-2K activation

#### Autophosphorylation at T348

Given that autophosphorylation at T348 represents a key step in eEF-2K activation, we first examined the ability of the individual CaM lobes to stimulate eEF-2K activity towards this site. We utilized constructs encoding individual lobes of CaM established by Sorensen and Shea (25), denoted CaM_N_ (residues 1-80) or CaM_C_ (residues 76-148). We incubated unphosphorylated eEF-2K (26) with ATP in the presence of 50 µM Ca^2+^ in the absence of CaM or 1 µM full-length wild-type CaM (CaM_WT_), CaM_N_, or CaM_C_. We quenched the reactions at various time points over an hour and monitored T348 phosphorylation by western blotting (Fig. 2A). As expected from previous studies that reported rapid T348 autophosphorylation with CaM_WT_ (*t*_1/2_ = 0.33 sec) (27), the reaction with CaM_WT_ was complete before the first time point was taken at 300 seconds. Autophosphorylation in the presence of CaM_C_ was similarly complete at the first time point. Conversely, in the presence of 1 or 10 µM CaM_N_, there was no significant increase in the rate of phosphate incorporation compared to that in the absence of CaM (Fig. S2). These data indicate that the isolated CaM_N_ cannot efficiently drive the activating autophosphorylation of eEF-2K under saturating Ca^2+^ and physiological concentrations of CaM.

**Fig. 2.**
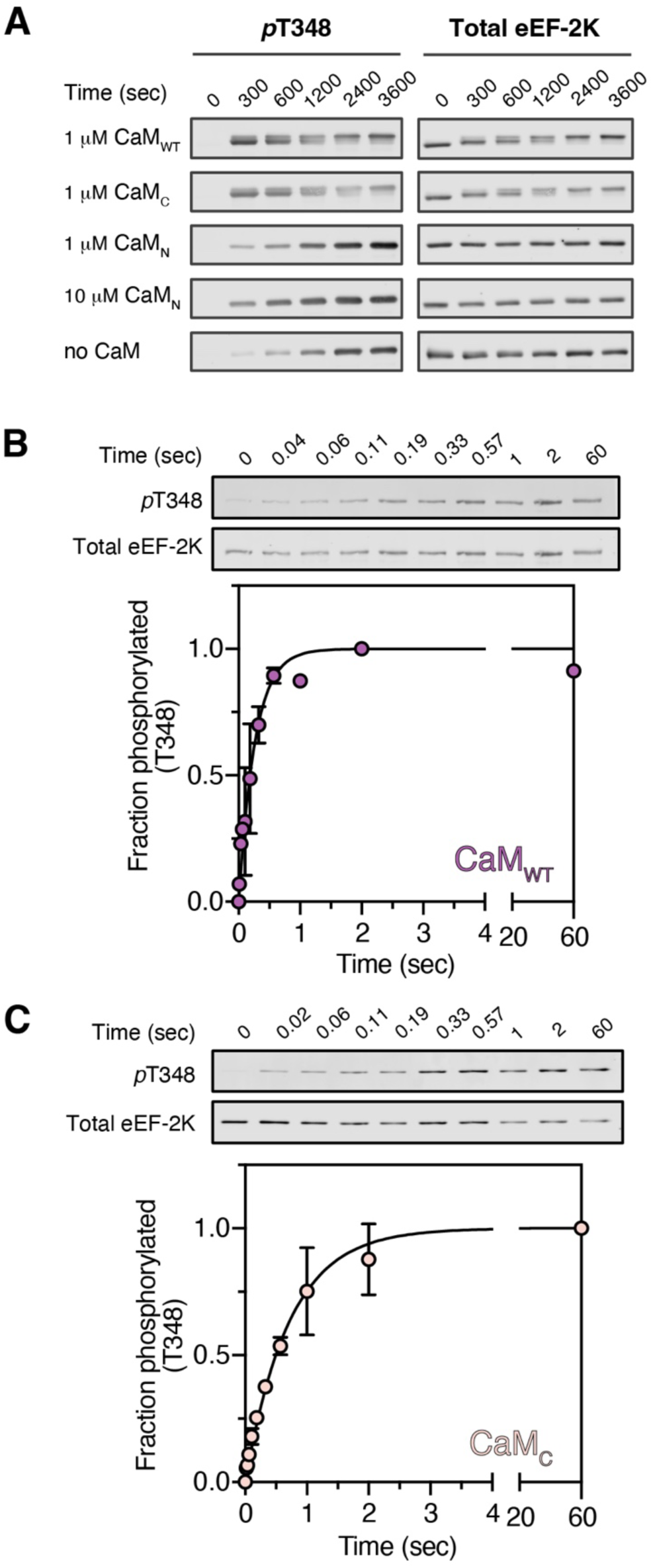
CaM_C_ stimulates eEF-2K autophosphorylation. **(A)** eEF-2K (300 nM) activity was measured against its primary autophosphorylation site, T348. The enzyme was incubated with CaM_WT_ (1 µM), CaM_C_ (1 µM), CaM_N_ (1 or 10 µM), or no CaM, and 50 µM free Ca^2+^ before initiating the reaction with 1 mM Mg^2+^•ATP. The reaction was quenched at various time points by adding 2.3 volumes of hot SDS-loading buffer. 150 ng of eEF-2K was loaded onto a gel, and *p*T348 and total eEF-2K were detected by western blotting. (**B-C**) Rapid quench flow was utilized to measure the rate of T348 autophosphorylation in the presence of CaM_WT_ or CaM_C_. eEF-2K (200 nM) was pre-incubated with (**B**) 2 µM CaM_WT_ or (**C**) 2 µM CaM_C_ in the presence of 50 µM free Ca^2+^, then rapidly mixed (2 ms) with 1 mM Mg^2+^•ATP. The reaction was quenched at various times with 200 mM KCl, 50 mM EDTA, and 10 mM EGTA. Western blotting was used to detect phosphorylation on T348, and phosphate incorporation was recorded as the fraction of their maximal control values (2 or 60 sec for CaM_WT_ or CaM_C_, respectively). The experimental data, shown as circles representing the mean with standard deviation (n = 2), were fit to Eq. 1 to obtain best-fit values of 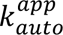 ; CaM_WT_ = 3.8 ± 0.38 s^-1^ and CaM_C_ = 1.4 ± 0.09 s^-1^.

We obtained the corresponding rates through rapid quench measurements to quantify T348 autophosphorylation stimulated by CaM_WT_ and CaM_C_. eEF-2K was pre-incubated with 2 µM CaM_WT_ or CaM_C_ and 50 µM free Ca^2+^, then rapidly mixed (∼2 ms) with saturating Mg^2+^•ATP. Reactions were quenched at various times and quantified by western blotting for *p*T348 and total eEF-2K (Fig. 2B, C). The autophosphorylation rates, (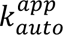) obtained by fits to Eq. 1 (see Experimental Procedures) were found to be similar for CaM_WT_ (3.8 ± 0.4 s^-1^) and CaM_C_ (1.4 ± 0.1 s^-1^) (Table 1).

**Table 1.**
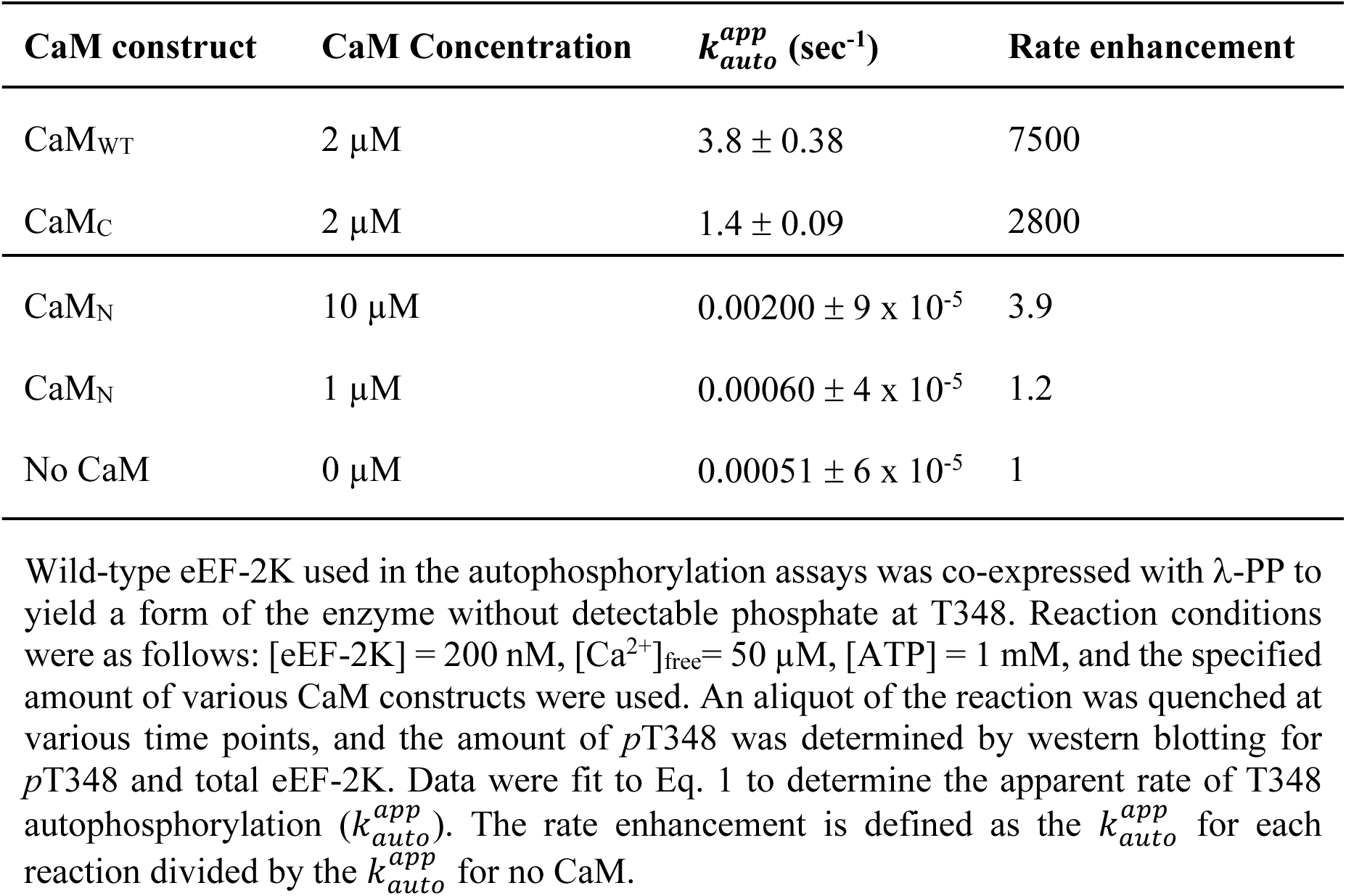
Rate of T348 autophosphorylation by eEF-2K.

| CaM construct | CaM Concentration | $k_{auto}^{app}$ (sec<sup>-1</sup>) | Rate enhancement |
| --- | --- | --- | --- |
| CaM<sub>WT</sub> | 2 μM | $3.8 \pm 0.38$ | 7500 |
| CaM<sub>C</sub> | 2 μM | $1.4 \pm 0.09$ | 2800 |
| CaM<sub>N</sub> | 10 μM | $0.00200 \pm 9 \times 10^{-5}$ | 3.9 |
| CaM<sub>N</sub> | 1 μM | $0.00060 \pm 4 \times 10^{-5}$ | 1.2 |
| No CaM | 0 μM | $0.00051 \pm 6 \times 10^{-5}$ | 1 |

Wild-type eEF-2K used in the autophosphorylation assays was co-expressed with $\lambda$ -PP to yield a form of the enzyme without detectable phosphate at T348. Reaction conditions were as follows: [eEF-2K] = 200 nM, [Ca<sup>2+</sup>]<sub>free</sub> = 50 μM, [ATP] = 1 mM, and the specified amount of various CaM constructs were used. An aliquot of the reaction was quenched at various time points, and the amount of *p*T348 was determined by western blotting for *p*T348 and total eEF-2K. Data were fit to Eq. 1 to determine the apparent rate of T348 autophosphorylation ( $k_{auto}^{app}$ ). The rate enhancement is defined as the $k_{auto}^{app}$ for each reaction divided by the $k_{auto}^{app}$ for no CaM.

#### Substrate phosphorylation

CaM stimulates eEF-2K’s activity towards substrate in a dose-dependent manner. The phosphorylation status of T348 kinetically influences substrate phosphorylation. Therefore, to elucidate the effect of CaM_C_ on eEF-2K’s activity toward a substrate, we utilized T348-phosphorylated eEF-2K (*p*eEF-2K) to measure its steady-state activity against a peptide substrate (PepS) under varying concentrations of CaM_WT_ or CaM_C_ in the presence of 50 µM free Ca^2+^ (Fig. 3A). The corresponding dose-response of eEF-2K activity to the concentration of CaM fitted to Eq. 2 (see Experimental Procedures) yielded similar 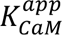 values (the concentration of CaM at half-maximal activity) for CaM_C_ (85 ± 10 nM) and CaM_WT_ (67 ± 7 nM). Further, the maximal observed activity (*k*_obs,max_) with saturating CaM_C_ (19.9 ± 0.7 s^-1^) and CaM_WT_ (20.0 ± 0.6 s^-1^) were similar in the two cases.

**Fig. 3.**
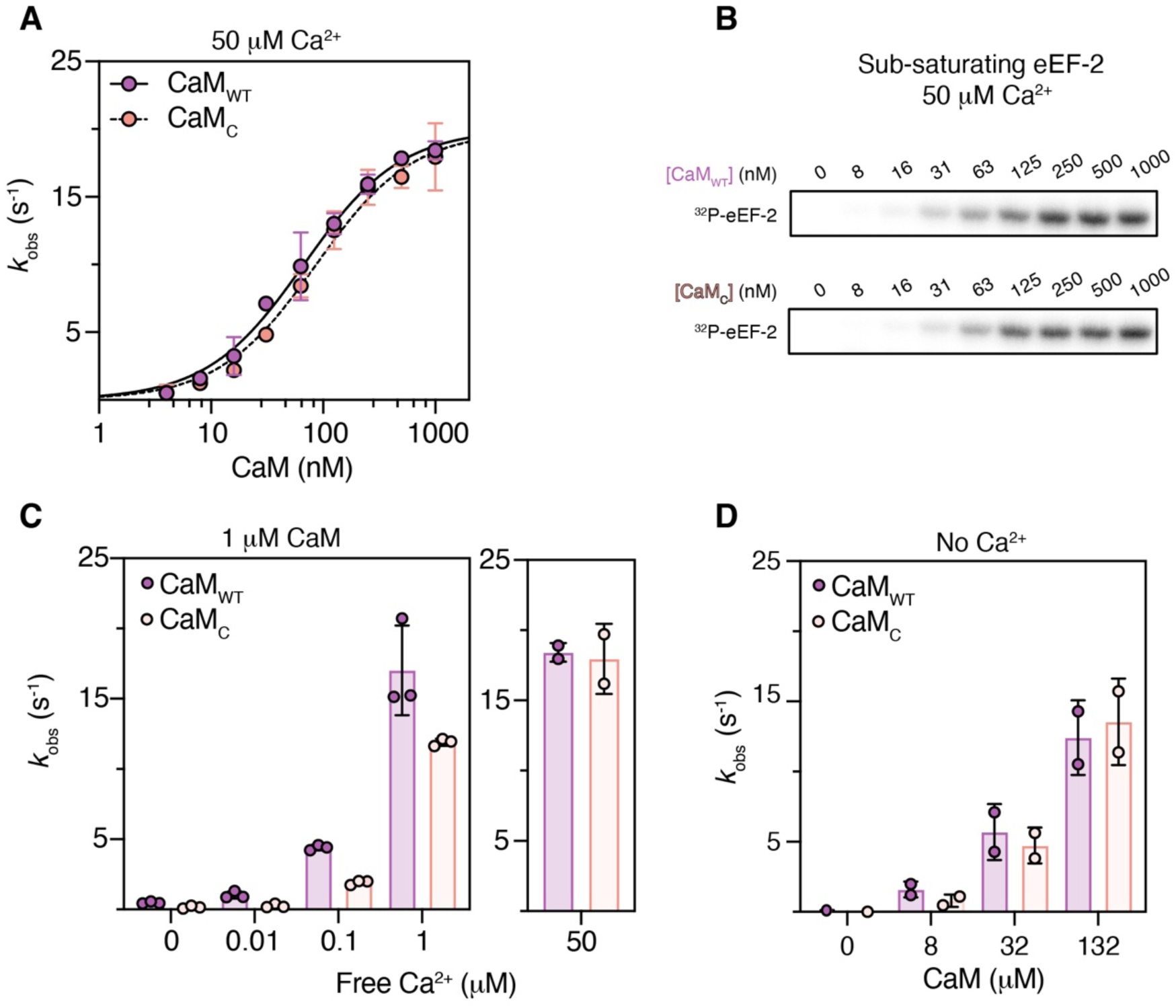
CaM_C_ binds and activates eEF-2K in a Ca^2+^-sensitive manner. **(A)** The dose-dependence of *p*eEF-2K (1 nM) activity on CaM_WT_ or CaM_C_ was measured against 150 µM peptide substrate (PepS) with Ca^2+^ (50 µM free) and 1 mM [γ-^32^P]-ATP. The *k*_obs_ values (mean with standard deviation, n = 2) were plotted against the concentration of the CaM construct, and data were fit to (Eq. 2) to obtain best-fit values for *k*_obs,max_ (CaM_WT_ = 20 ± 0.6 s^-1^; CaM_C_ = 20 ± 0.7 s^-1^) and 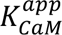 (CaM_WT_ = 67 ± 7 nM; CaM_C_ = 85 ± 10 nM). **(B)** The activity of 0.5 nM eEF-2K was measured against 5 µM of yeast eEF-2 in the presence of Ca^2+^ (50 µM free) with 1 mM [γ-^32^P]-ATP and varied CaM_WT_ or CaM_C_. Samples were quenched in addition to a hot SDS-loading buffer and then analyzed using SDS-PAGE gels. A phosphorimager visualized the incorporation of ^32^P into eEF-2. Samples were extracted from the gel and measured by scintillation counting. The *k*_obs_ at each CaM concentration were plotted against the CaM concentration, and data were fit to (Eq. 2) to obtain best-fit values for *k*_obs_,_max_ (CaM_WT_= 6.1 ± 1.5 s^-1^; CaM_C_ = 5.6 ± 1.4 s^-1^) and 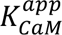 (CaM_WT_= 121 ± 41 s^-1^; CaM_C_ = 156 ± 64 s^-1^). Also see Fig. S3. (**C**) The activity of 2 nM eEF-2K was measured against 150 µM PepS with 1 mM [γ-^32^P]-ATP in the presence of 1000 nM CaM_WT_ or CaM_C_ with varying concentrations of free Ca^2+^ (0-1000 nM). The *k*_obs_ values obtained at various Ca^2+^ concentrations are indicated. On the right, Data from (A), 50 µM free Ca^2+^ and 1000 nM CaM_WT_ or CaM_C_ are re-plotted for comparison. **(D)** The activity of 2 nM eEF-2K with varying concentrations of CaM_WT_ or CaM_C_ in the absence of Ca^2+^ was measured against 150 µM PepS with 1 mM [γ-^32^P]-ATP; the corresponding *k*_obs_ values are shown.

To assess whether CaM_C_ has a similar stimulatory effect relative to CaM_WT_ towards eEF-2, as noted for the peptide substrate, we measured the activity of *p*eEF-2K against yeast eEF-2 using varying concentrations of CaM_WT_ or CaM_C_ in the presence of Ca^2+^ (Fig. 3B). Use of a sub-saturating concentration of eEF-2 in this assay ensures that a variation in *k*_obs,max_ would be detected as a change in catalytic efficiency. We found that the 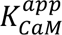 and *k*_obs,max_ values obtained for CaM_C_ were not substantially different from those for CaM_WT_ (Fig. S3).

Although our data suggest that under the tested conditions, CaM binding and eEF-2K activation are unaffected by the absence of CaM_N_, it should be noted that kinase activity was measured with a Ca^2+^: Mg^2+^ ratio (1:200) that highly favors Mg^2+^ (similar to the cellular ratio under resting conditions). It is possible that under these conditions, Mg^2+^ binding to the N-lobe of CaM_WT_ may mask its contributions to complex formation (18, 28). To exclude this scenario, we measured the 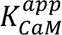 and *k*_obs,max_ of *p*eEF-2K against PepS with CaM_WT_ and CaM_C_ using an altered Ca^2+^: Mg^2^ (1:10), without any apparent effect on the 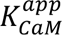 for either CaM construct (Fig. S4).

#### Calcium sensitivity

Ca^2+^ enhances the 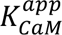 of *p*eEF-2K toward a peptide substrate by approximately 300-fold (26, 27). To investigate whether a similar effect exists for CaM_C_, we measured *p*eEF-2K activity against PepS with a fixed concentration of CaM_WT_ or CaM_C_ at various concentrations of free-Ca^2+^ (Fig. 3C). Previous studies have reported that the Ca^2+^ affinity of the CaM_C_ construct is slightly reduced compared to CaM_WT_ (25). It was, therefore, not surprising that 1 nM *p*eEF-2K with 1 µM CaM_C_ displayed full activity when stimulated with a high concentration of Ca^2+^ (50 µM), but activity at reduced Ca^2+^ concentrations was lower for CaM_C_ compared to CaM_WT_. Overall, the activity of *p*eEF-2K with CaM_C_ or CaM_WT_ increased with increasing Ca^2+^. Given that Ca^2+^ does not appear to directly simulate eEF-2K in the absence of CaM (26), these data suggest that Ca^2+^ bound to CaM_C_ enhances eEF-2K activity by promoting complex formation. Notably, CaM_WT_ and CaM_C_ show comparable abilities to activate eEF-2K in the absence of Ca^2+^ (Fig. 3D).

### Fusing CaM_C_ to eEF-2K leads to a constitutively active kinase

#### Expression, purification, and characterization of a chimeric construct

Based on the evidence above, CaM_C_ can fully activate eEF-2K. The primary role of Ca^2+^ appears to be to enhance the affinity of CaM_C_ for eEF-2K, leading to its activation. Therefore, it can be expected that enhancing the local concentration of CaM, specifically of CaM_C_, will enable kinase activation with reduced reliance on stimulatory inputs such as Ca^2+^. Leveraging our previous structural information (18, 22, 23) and our knowledge of eEF-2K activation by CaM_C_ (described above), we developed a chimeric construct (Fig. 4A) in which CaM<u>_C_</u> is <u>Li</u>nked to <u>N</u>-truncated eEF-2<u>K</u> (C-LiNK). This construct comprises CaM_C_ (76–148) fused to N-terminally truncated eEF-2K (71–725) using a linker comprising two glycine residues. This linkage places CaM_C_ immediately upstream of the CaM-targeting motif (CTM, see Fig. S1), which is key for the CaM/eEF-2K interaction (18, 29). We have previously shown that the disordered N-terminus of eEF-2K is dispensable for its activation by CaM (24). C-LiNK was expressed (Fig. S5A, B) and purified similarly to wild-type eEF-2K (30), and its monomeric state was confirmed by multi-angle light scattering (MALS) that produced a single peak at a molecular mass of ∼82.4 KDa (Fig. 4B).

**Fig. 4.**
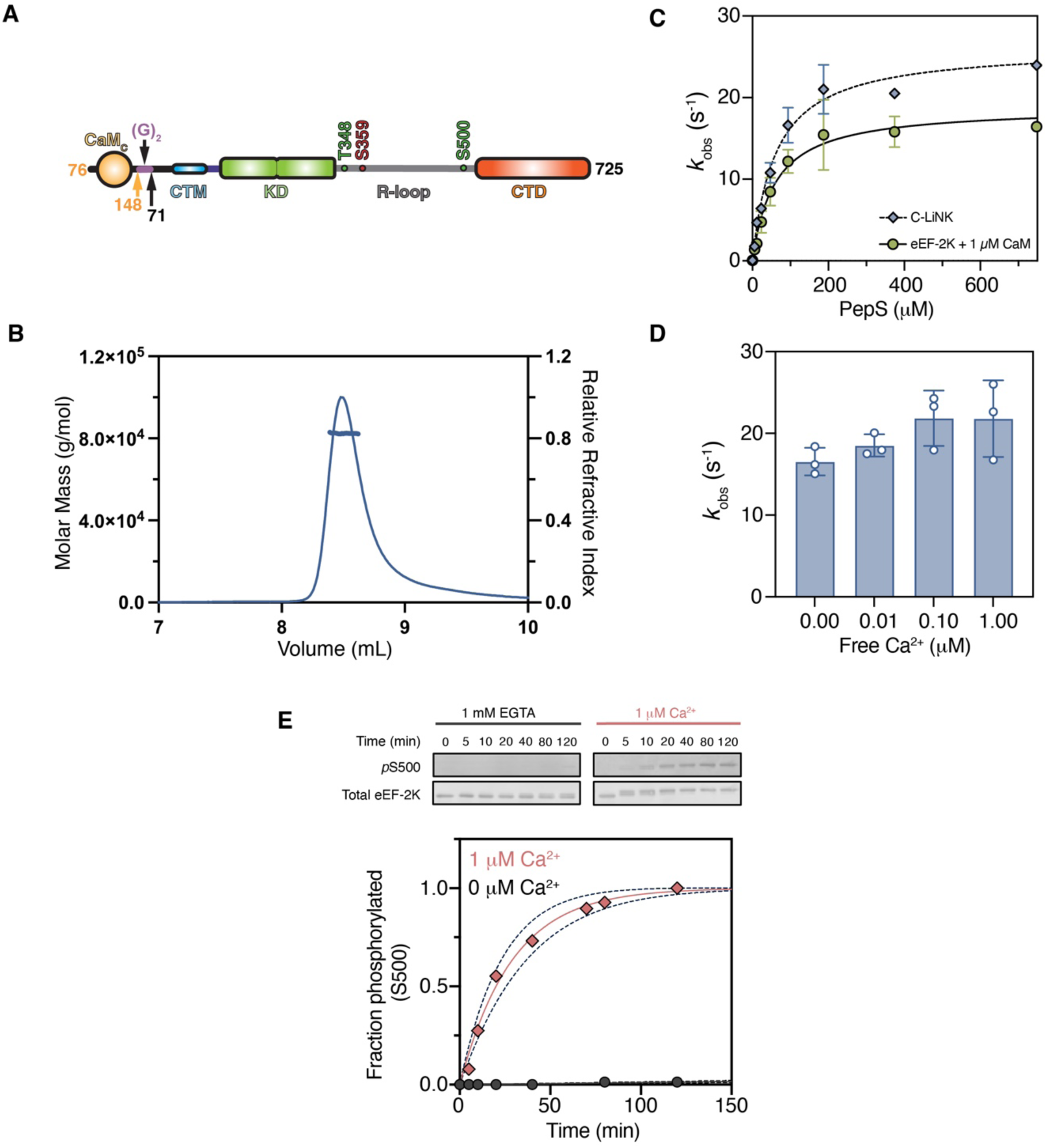
In vitro characterization of C-LiNK. **(A)** A schematic representation of the C-LiNK construct in which CaM_C_ (CaM residues 76-148) is connected via two glycine residues to an N-terminally truncated eEF-2K (71–725). The various key elements of eEF-2K, including the CaM-targeting motif (CTM), the α−kinase domain (KD), the regulatory loop (R-loop) with activating (T348, S500) and inhibiting (S359) phosphorylation sites are indicated schematically. **(B)** Multi-angle light scattering (MALS) analysis of purified C-LiNK is shown. The curve represents the sample’s refractive index, while the horizontal data points represent the estimated molar mass across the peak. C-LiNK is found to be monomeric, with a molar mass of ∼82.4 kDa. **(C)** The activity dependence of 1 nM wild-type eEF-2K (with 1 µM CaM) or C-LiNK (with 0 µM added CaM) on PepS concentration was measured using 1 mM [γ-^32^P]-ATP in the presence of 50 µM free Ca^2+^. The *k*_obs_ was plotted against PepS concentration and fit to Eq. 3 to obtain best-fit values for 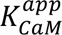(eEF-2K = 19 ± 1 sec^-1^; C-LiNK = 26 ± 1 sec^-1^) and *K*_m_ (eEF-2K = 59 ± 12; C-LiNK= 61 ± 9 µM). **(D)** The activity of 1 nM C-LiNK was measured against 150 µM PepS using 1 mM [γ-^32^P]-ATP, 10 mM free Mg^2+^, 1 mM EGTA, and various amounts of CaCl_2_. The open circles indicate experimental data points (n=3). **(E)** Measurement of C-LiNK (300 nM) activity towards the secondary autophosphorylation site, S500. Reactions were incubated with 0 or 1 µM free Ca^2+^ at 30 °C before initiating with 1 mM ATP. Samples were subject to western blotting to detect total enzyme and *p*S500 (ECM Biosciences) levels; 0.125 µg of protein was loaded in each case. Signals for *p*S500 and total C-LiNK were quantified, and *p*S500 levels were corrected for their corresponding total C-LiNK signals, then converted to fraction phosphorylated by normalizing data to the 1 µM Ca^2+^ 120 min sample. The experimental data, shown as circles representing the mean (n = 2) and standard deviation, were fit to Eq. 1 (lower panel). The solid line indicates the best fit through the data. The best fit 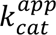 for 0 µM free Ca^2+^ could not be ascertained (experimental data shown as black filled circles); 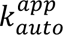 for 1 µM free Ca^2+^ (experimental data shown as red filled circles) was 0.00057 ± 0.6 ξ 10^-4^ sec^-1^ (*t*_1/2_ ∼20 min).

C-LiNK was co-expressed with lambda phosphatase (λ-PP), a procedure used to express WT eEF-2K devoid of phosphorylation (26). However, despite this procedure, phosphorylation on T348 (we use native eEF-2K numbering in all cases) was detected for C-LiNK by western blot, and further incubation of purified C-LiNK with ATP did not increase the intensity of the *p*T348 signal (Fig. S5C).

eEF-2K can undergo several secondary autophosphorylations, many highly dependent on CaM (26, 27, 31). Given the apparent high basal activity of C-LiNK, we utilized a bottom-up tandem mass spectrometry (MS) approach to confirm the phosphorylation status of the expressed protein. C-LiNK was proteolytically digested by trypsin and then analyzed using LC-MS/MS. The resulting MS/MS spectra accounted for 99% sequence coverage, and phosphorylation phosphorylation was detected at residues T348 (>99%) and S445 (1%) (Fig. S6). Fragment ion analysis, including assignment of b- and y-ion series with sub-10 ppm mass accuracy, supported high-confidence peptide identification and phosphorylation site assignment (Table S1 and S2). Next, we confirmed the ability of C-LiNK to autophosphorylate on T348, S445, and S500 by western blotting using specific antibodies after incubating the purified protein with Mg^2+^•ATP for varying periods. This confirmed that C-LiNK can undergo autophosphorylation at these residues, similar to CaM-stimulated wild-type eEF-2K CaM (Fig. S5C-E).

#### Ca^2+^ dependence of C-LiNK activity

To assess C-LiNK activity relative to CaM-stimulated WT eEF-2K, we compared the enzymatic activities of C-LiNK or *p*eEF-2K (using 1 µM CaM) against PepS in the presence of saturating Ca^2+^. Fits to Eq. 3 (see Experimental Procedures; Fig. 4C) indicate that C-LiNK is fully active against the peptide substrate (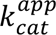 = 26 ± 1 s^-1^ for C-LiNK versus 19 ± 1 s^-1^ for *p*eEF-2K), with similar catalytic efficiency (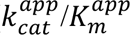 = 0.43 ± 0.05 µM^-1^ s^-1^ for C-LiNK, 0.32 ± 0.05 µM^-1^ s^-1^ for *p*eEF-2K) as wild-type CaM-stimulated eEF-2K. The activity of C-LiNK towards the peptide substrate was independent of added Ca^2+^ (0-1000 nM of free Ca^2+^; Fig. 4D). Extended dialysis against a Ca^2+^-free buffer (1 mM EGTA for 19 hours) before measuring the catalytic activity had no impact on the results (Fig. S7).

It has previously been shown that Ca^2+^-CaM is necessary for efficient autophosphorylation at the activating site S500, as is prior phosphorylation at T348 (27). To investigate this further using C-LiNK, which is fully phosphorylated at T348, we incubated C-LiNK with Mg^2+^-ATP in the presence of 1 µM free Ca^2+^ and monitored the appearance of *p*S500 over time (Fig. 4E). The data revealed that C-LiNK has a similar rate of S500 autophosphorylation (*t*_1/2_ ∼20 min) as WT eEF-2K in the presence of saturating Ca^2+^-CaM (26). However, this rate is markedly reduced in the presence of 1 mM EGTA without added Ca^2+^ (Fig. 4E), suggesting that the autophosphorylation of S500 remains reliant on Ca^2+^ as in the case of wild-type eEF-2K.

#### C-LiNK displays a similar functional core as the CaM•peEF-2K complex

As noted above, C-LiNK is constitutively active and retains all the properties of wild-type eEF-2K upon its activation by CaM (and CaM_C_). One could, therefore, expect that the overall conformation of its functional core, key CaM-recognition modules, and catalytic residues would not be substantially different from that seen in the structures of the CaM•*p*eEF-2K_TR_ complex. To confirm this hypothesis, we solved the structure of C-LiNK_TR_ (Fig. 5A) by x-ray crystallography (see Table 2 for details of data collection, refinement, and structure statistics). The CaM_C_ and eEF-2K structural modules maintain their overall conformations in the context of C-LiNK_TR_ as seen in the structures CaM•*p*eEF-2K_TR_ complex solved in various states (Fig. S8). The average deviation over the CaM_C_ and eEF-2K_TR_ modules was 0.5 ± 0.1 Å (533 ± 12 trimmed residues over 5 distinct structures). The 7SHQ structure that shows a greater degree of closure between the kinase domain and CTD (Fig. S8) was excluded from the average.

**Fig. 5.**
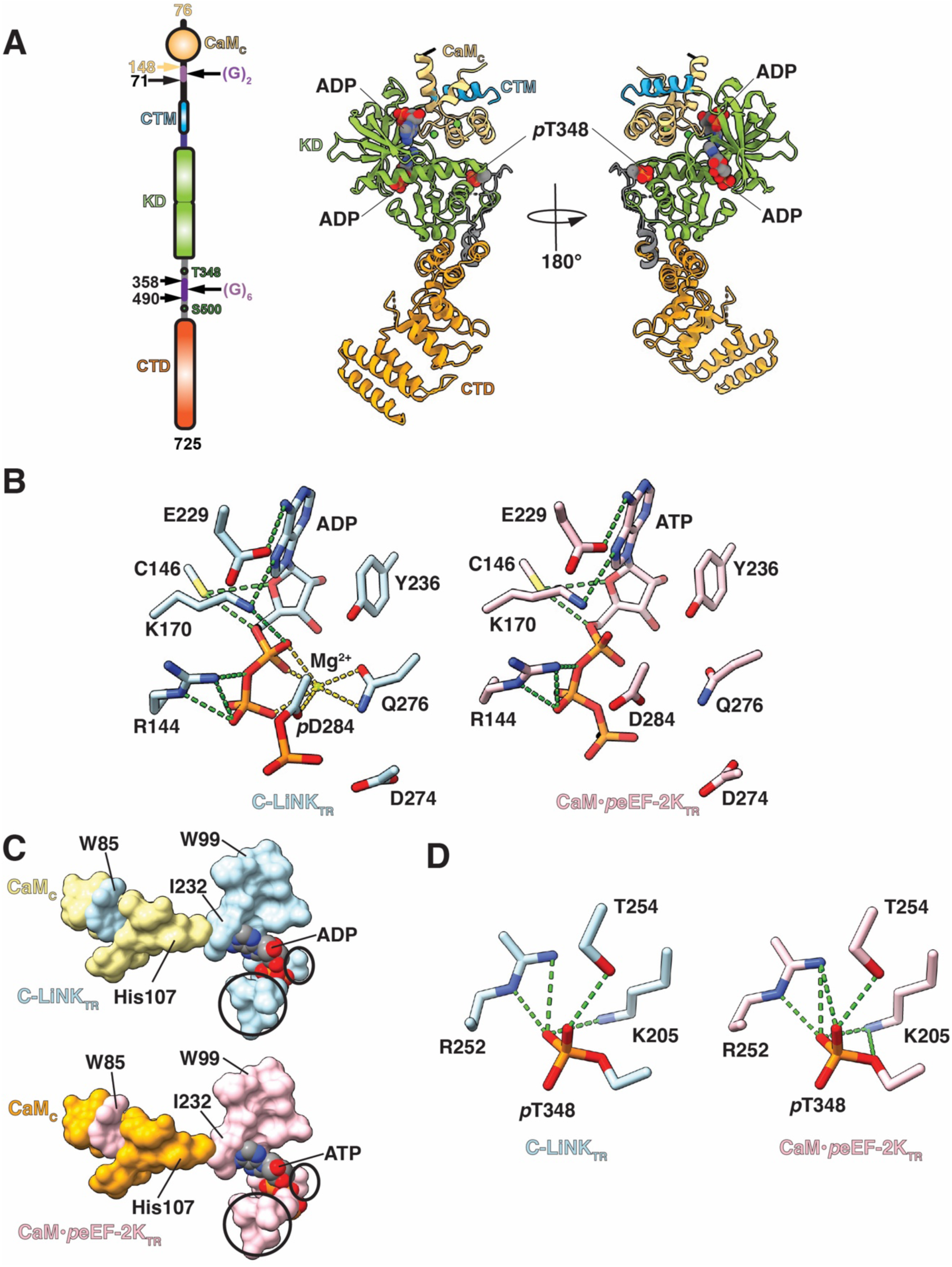
Structure of C-LiNK_TR_. **(A)** The organization of the C-LiNK_TR_ construct is shown schematically on the left panel. This construct is derived from C-LiNK (Fig. 4A) with an R-loop segment comprising residues 359-489 replaced by 6 glycines in analogy to eEF-2K_TR_. The structure of C-LiNK_TR_ is shown on the right panel, with the key structural modules indicated and colored as on the left panel. Two ADP molecules, one bound at the catalytic site and a second bound to the interface between the CaM_C_ module and the N-lobe of the KD, are shown as spheres. The phosphorylated T348 (*p*T348) is also indicated. **(B)** Conformation of key catalytic site elements in C-LiNK (left) and the CaM•*p*eEF-2K_TR_ complex (PDB: 8FNY, right) show no significant variation in conformation. D284 in the C-LiNK structure is phosphorylated and contains a bound Mg^2+^ ion. Hydrogen bonds are indicated by the green dashed lines in all cases; the gold dashed lines denote heteroatoms with 3.2 Å of the metal center. **(C)** The activation spine that links CaM_C_ to the kinase catalytic site through the bound nucleotide is fully formed in C-LiNK. The geometry of the spine in C-LiNK (top panel; eEF-2K modules in light blue, CaM_C_ in yellow) is identical to that seen in the structures of the CaM•eEF-2K_TR_ complex (bottom panel; a representative heterodimeric complex, PDB: 8FNY; eEF-2K in pink, CaM_C_ in orange). Key spine residues are labeled (3-letter code for the CaM residue), the nucleotide bound to the catalytic site is shown in both cases, and the active site is circled. **(D)** The coordination of *p*T348 at the phosphate-binding pocket in C-LiNK (left) and the 8FNY structure (right) is identical.

**Table 2.** Data collection and structure refinement statistics^a^.

|  |  |
| --- | --- |
| Wavelength (Å) | 0.92022 |
| Space Group | P12<sub>1</sub>1 |
| Unit cell a, b, c (Å) | 64.766 59.736 88.766 |
| Unit cell $\alpha$, $\beta$, $\gamma$ (°) | 90.0 98.497 90.0 |
| Resolution range for data processing (Å) | 49.39 - 2.271 (2.352 - 2.271) |
| Total number of observations | 124377 (6027) |
| Total number of unique observations | 17519 (876) |
| Mean(I)/$\sigma$(I) | 6.6 (1.6) |
| Completeness (spherical) | 55.9 (9.6) |
| Completeness (ellipsoidal)<sup>b</sup> | 91.7 (63) |
| Multiplicity | 7.1 (6.9) |
| CC<sub>1/2</sub> | 0.994 (0.661) |
| $R_{\text{merge}}$ (all I<sup>+</sup> & I<sup>-</sup>) | 0.218 (1.228) |
| $R_{\text{meas}}$ (all I<sup>+</sup> & I<sup>-</sup>) | 0.235 (1.328) |
| $R_{\text{pim}}$ (all I<sup>+</sup> & I<sup>-</sup>) | 0.088 (0.503) |
| Reflections used in refinement | 17506 (73) |
| Reflections used for $R_{\text{free}}$ | 858 (2) |
| $R_{\text{work}}$ | 0.1996 (0.3123) |
| $R_{\text{free}}$ | 0.2559 (0.5003) |
| Number of non-hydrogen atoms | 4636 |
| Macromolecules | 4421 |
| Ligands | 84 |
| Solvent | 155 |
| Protein residues | 566 |
| RMS (bonds) | 0.011 |
| RMS (angles) | 0.71 |
| Ramachandran favored (%) | 96.0 |
| Ramachandran allowed (%) | 3.64 |
| Ramachandran outliers (%) | 0.36 |
| Rotamer outliers (%) | 3.04 |
| Clash Score | 14.65 |
| Average B-factor (Å<sup>2</sup>) | 29.32 |
| Macromolecules | 29.51 |
| Ligands | 32.06 |
| Solvent | 22.98 |
| PDB accession code | 9OCW |

<sup>a</sup> Statistics for the highest-resolution shell are shown in parentheses.
<sup>b</sup> Diffraction limits & principal axes of ellipsoid fitted to diffraction cut-off surface
3.411 Å; 0.9481 0.0000 0.3179; 0.970a\* + 0.244c\*
2.498 Å; 0.0000 1.0000 0.0000; b\*
2.253 Å; -0.3179 0.0000 0.9481; -0.229a\* + 0.973 c\*

The catalytic site contains a bound ADP molecule and a single Mg^2+^ ion, with the orientations of key sidechains of the catalytic residues being essentially unchanged from that seen in the structures of the CaM•*p*eEF-2K_TR_ with a nucleotide (ATP) bound to the catalytic site (Fig. 5B). We have previously noted the presence of an “activation spine (A-spine)” that links the CTM engaged CaM_C_ through a conserved regulatory element (RE) and the bound nucleotide to the kinase active site (18). Not surprisingly, the A-spine is fully formed in C-LiNK, with the constituent residues in similar configurations to the nucleotide-bound form of the CaM•eEF-2K_TR_ complex (Fig. 5C).

Surprisingly, even though a fully dephosphorylated C-LiNK was used for crystallization, T348 was found to be phosphorylated and engaged to the phosphate-binding pocket similar to that seen in the structures of the CaM•eEF-2K_TR_ complex (Fig. 5D). Additionally, D284, at the catalytic site was also phosphorylated (Fig. 5B) as seen previously (in preparation). Notably, the equivalent residue (D766) in the kinase domain of the homologous *Dictyostelium* myosin heavy chain kinase A (*ds*MHCK-A) has also been seen in the phosphorylated form in multiple crystal structures (32, 33). We surmised that an ATP contaminant in the ADP used in the crystallization buffer could be the origin of the phosphates, with the phospho-transfer occurring *in crystallo*. Given that the level of ATP contamination of our ADP stock was ∼2% by NMR analysis, this seems to be a somewhat unlikely scenario.

To test the origin of these multiple phosphorylations despite the apparent lack of ATP, C-LiNK_TR_ was incubated with Ca^2+^ and ADP (from the stock solution used in crystallization) without Mg^2+^. The resulting molecular mass, measured by ESI-QTOF-mass spectrometry, was consistent with that of the unmodified protein. Adding Mg^2+^ resulted in a shift of ∼80 Da, indicating a single phosphorylation event (Fig. S9) and suggesting that C-LiNK can perhaps utilize ADP to drive phospho-transfer. It has been noted that *ds*MHCK-A can also use ADP to drive substrate phosphorylation (32). This curious property will be explored elsewhere.

#### Expression of C-LiNK renders cellular peEF-2 levels insensitive to stimuli

Our *in vitro* analysis of C-LiNK established that it is fully active and exhibits properties similar to CaM-bound wild-type eEF-2K. To assess the cellular properties of C-LiNK, we transfected MCF10A^eEF-2K-/-^ cells with pcDNA3 encoding tagless C-LiNK or wild-type eEF-2K. Given that previous studies have suggested that high levels of cellular eEF-2K activity correlate with increased degradation (27), it was not surprising that cells displayed lower levels of C-LiNK protein relative to WT eEF-2K (Fig. 6A). Mutation of the D284 (described above) to alanine results in a loss of catalytic activity in eEF-2K (27). Transfection of the D284A mutant of eEF-2K and C-LiNK rescued protein levels in MCF10A^eEF-2K-/-^ cells, confirming that the low levels of C-LiNK protein (and of wild-type eEF-2K) are indeed correlated with its activity (Fig. S10).

**Fig. 6.**
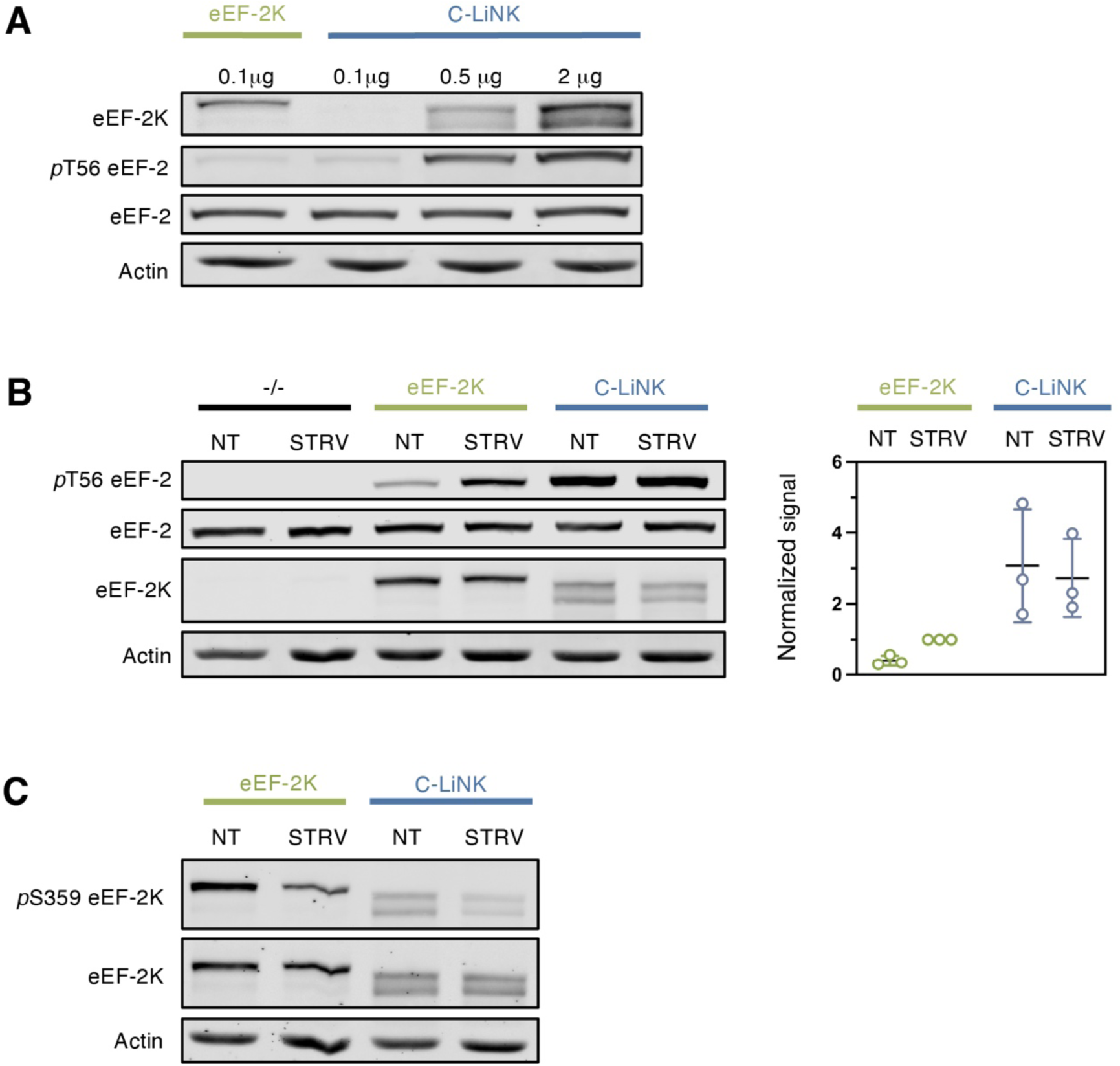
C-LiNK activity in MCF10A^eEF-2K-/-^ cells resists inhibitory signaling. **(A)** MCF10A^eEF-2K-/-^ cells were transfected with the indicated amount of vector encoding either wild-type eEF-2K or C-LiNK. After 16 hours, cells were lysed and analyzed by western blotting using specific antibodies for eEF-2K, eEF2, *p*eEF-2, and actin (loading control). **(B)** MCF10A^eEF-2K-/-^ cells transfected with a vector encoding no gene, WT eEF-2K, or C-LiNK. After 16 hours, cells were treated for 2 hours with complete media (NT) or starved with DPBS (STRV). The experiment was performed in triplicate, and a representative western blot is shown. The graph shows the *p*eEF-2 signal corrected by the total eEF-2 signal, then normalized as the fraction of the signal for wild-type eEF-2K STRV. **(C)** Wild-type eEF-2K or C-LiNK expressing cells were starved or given fresh media, and the abundance of the inhibitory phosphorylation on S359 was assessed via western blotting.

To account for the reduced cellular levels of C-LiNK relative to wild-type eEF-2K, we titrated MCF10A^eEF-2K-/-^ cells with increasing C-LiNK pcDNA3 vector and immunoblotted for eEF-2K. A 5-fold increase in the concentration of C-LiNK pcDNA3 relative to wild-type eEF-2K pcDNA3 was sufficient to produce similar protein levels and was used for further cellular experiments. Interestingly, the eEF-2K antibody recognizes C-LiNK as two distinct bands. A similar banding pattern is observed for eEF-2K *in vitro* when incubated with CaM and ATP (for example, see Figs. 2A and 4E). We suspect that C-LiNK can undergo significant autophosphorylation in cells, which may be the origin of these additional bands. The fact that the kinase-dead (D284A) mutant is recognized as a single band is consistent with this scenario (Fig. S10).

Cellular eEF-2K activity has been reported to be highly nutrient-sensitive. Nutrients and growth factors suppress eEF-2K activity through mTOR-mediated phosphorylation of residues within the R-loop (e.g., on S359) (34). Removing nutrients (starvation) from cellular media decreases these inhibitory phosphorylation events and enhances eEF-2 phosphorylation (34). MCF10A^eEF-2K-/-^ cells expressing wild-type eEF-2K in nutrient-rich (non-treated, NT) media showed lower levels of *p*eEF-2 than cells subjected to starvation (STRV, Fig. 6B). The activity of C-LiNK, as measured by *p*eEF2 levels, was higher than wild-type eEF-2K in both NT and STRV conditions and displayed no nutrient sensitivity. We also observed a reduction in S359 phosphorylation for both constructs in starved (STRV) MCF10A^eEF-2K-/-^ cells as compared to NT cells (Fig. 6C). These data indicate that while levels of inhibitory phosphorylation can be altered on C-LiNK by regulatory inputs, as in WT eEF-2K, its activity isis insensitive to these modifications.

Finally, to confirm that endogenous CaM did not drive C-LiNK activation in the cellular context, we tested the ability of C-LiNK to interact with CaM. We found that fluorescently labeledCaM_WT_ (I-CaM) cannot engage C-LiNK, even when present in a 20-fold molar excess. In contrast, WT eEF-2K shows robust binding to I-CaM (Fig. S11). Thus, it may be concluded that the cellular activity of C-LiNK results from activation in *cis* through its N-terminal CaM_C_ rather than through its interaction with cellular CaM.

## Discussion

CaM’s role as a Ca^2+^ sensor necessitates adaptive conformational changes, which enable it to bind and activate numerous target proteins with varying Ca^2+^ sensitivities (35, 36). Regulation of most CaM targets, including other CaM-dependent kinases, generally requires interaction with both lobes of CaM (37–39). Although rare, examples in the literature describe targets that can be stimulated to varying degrees by individual lobes of CaM (40). For CaM-regulated kinases, the stimulatory dominance of one lobe of CaM over the other has been noted, e.g., the isolated CaM_N_ appears to partially activate CaMKII in contrast to CaM_C,_ which cannot (41). Nevertheless, eEF-2K is unique in that maximal activation can be stimulated by the isolated CaM_C_. This lobe also appears to drive the Ca^2+^-sensitivity of activation. CaM lobe-centric activity has been suggested to play a role in the temporal response of decoding diverse Ca^2+^ signals, leading to distinct downstream responses (42, 43). A similar role in interpreting and transducing specific Ca^2+^ fluxes to modulate protein synthesis may also be operative with respect to eEF-2K.

It has been suggested that eEF-2K activity is regulated by Ca^2+^ and post-translational modifications through modulation of its affinity for CaM (8). To investigate this further, we leveraged our structural understanding of the CaM/eEF-2K complex and our discovery of the capacity of CaM_C_ to activate eEF-2K to create a constitutively active kinase by genetically linking CaM_C_ to an N-terminally truncated eEF-2K (C-LiNK). It is to be noted that this contrasts with “conventional” CaM-regulated kinases, where deletion of an auto-inhibitory or pseudo-substrate segment leads to constitutive activation (44, 45). The Ca^2+^-dependent protein kinases (CDPKs) found in plants and protozoans are genetic fusions of CaMK and CaM-like domains. Given that the CDPKs are also activated through the “release of inhibition” mechanism operative in conventional CaM-regulated kinases, they may also be constitutively active by deleting the autoinhibitory domain (46–48). This makes C-LiNK a mechanistically unique species.

C-LiNK activity towards a peptide substrate appears independent of the presence of Ca^2+^. Indeed, extended incubation with the Ca^2+^-chelator EGTA did not affect the activity. We cannot fully discount the scenario that the fusion greatly enhances the affinity for Ca^2+^ within the fused CaM_C_ to such an extent that the ions cannot be removed through EGTA incubation. Indeed, a marked increase in Ca^2+^ affinity of CaM fused to a CaM-binding peptide has been noted previously (49). Native mass spectrometric analyses proved inconclusive in our case. Nevertheless, the CaM_C_ module within the C-LiNK construct can engage Ca^2+^, as evidenced by the structure of the C-LiNK_TR_ and the fact that Ca^2+^ is necessary to stimulate autophosphorylation at S500 in line with our previous observations for wild-type eEF-2K (27). However, the presence of additional Ca^2+^-binding sites elsewhere on eEF-2K, perhaps on the disordered R-loop, that may influence the latter event cannot be ruled out by the present studies.

C-LiNK appears fully active when transiently expressed in MCF10A^eEF-2K-/-^ cells under nutrient-rich and nutrient-deprived conditions. This contrasts with wild-type eEF-2K, whose nutrient deprivation significantly enhances activity. Under nutrient-rich conditions, the inhibitory phosphorylation (S359) status on C-LiNK mimicked that of wild-type eEF-2K. However, the activity of C-LiNK, as measured by *p*eEF2 levels, was sustained despite this suppressive phosphorylation, suggesting that C-LiNK is active under normally inhibitory conditions. This is consistent with the scenario that inhibitory phosphorylation downregulates eEF-2K activity by reducing CaM binding but has no effect once CaM is stably engaged to eEF-2K.

Although the stimulatory contribution of CaM_N_ was negligible under our conditions, it may have an alternative role that has not been explored here. It is possible that CaM_N,_ which is more sensitive to rapid Ca^2+^ transients, could play a role in eEF-2K activity upon induction of specific Ca^2+^ signals (43). It is also possible that CaM_N_ could play a greater role under the influence of specific post-translational modifications not explored here. Additional experiments are needed to assess the presence and magnitude of these effects.

## Experimental procedures

### Protein expression and purification

#### Calmodulin constructs

Recombinant CaM_N_ (1–80) was purchased from the University of Iowa Proteomics Facility. Recombinant CaM_WT_ and CaM_C_ (76–148) were expressed and purified as described previously (41, 50). Briefly, CaM_WT_ in pET-23 or CaM_C_ in T7-7 (a gift from Dr. Madeline Shea, U. of Iowa) (25) vectors were expressed in BL21 (DE3) pLysS *E. coli* cells. The cell pellets were resuspended in 50 mM Tris (pH 7.5) with 10 mM EDTA and protease inhibitors (TPCK, PMSF, and Benzamidine) and sonicated to lyse the cells. The lysate was centrifuged at 27,200 x g for 30 minutes at 4 °C, and the supernatant was collected. The CaM constructs were purified by heating the supernatant to 80 °C for 20 min, then centrifuging at 2,000 x g for 15 min at 22 °C and filtering the resulting supernatant. The sample was then applied to the HiPrep Phenyl FF 16/10 column and washed with 2 column volumes (CV) of purification buffer A [50 mM Tris (pH 7.5) with 1 M CaCl_2_] and eluted with 2 CV of purification buffer A containing 0.5 M NaCl or purification buffer B [50 mM Tris (pH 7.5) with 10 mM EGTA]. Then, the sample was passed through a HiPrep 26/10 desalting column (Amersham Biosciences), applied to a Mono-Q HR 10/10 (Amersham Biosciences), washed with 5 CV of 50 mM Tris (pH 7.5), and eluted over 20 CV with a gradient up to 250 mM KCl in 50 mM Tris (pH 7.5). The sample was dialyzed into 25 mM HEPES (pH 7.5) for storage.

#### eEF-2K constructs (WT, C-LiNK, and C-LiNK_TR_)

A pET32a plasmid containing the designed C-LiNK sequence was purchased from GenScript. Recombinant eEF-2K constructs were expressed and purified as previously described, with minor adjustments to the purification protocol (30). Constructs were expressed either alone or in combination with λ-phosphatase (pCDF-duet vector), as indicated by the presence or absence of the prefix ‘p’, respectively (e.g. ‘eEF-2K’ denotes co-expression with λ-phosphatase, while ‘*p*eEF-2K’ indicates expression without it). Cell lysis and Ni-NTA affinity chromatography were performed as previously described (30). Following the Ni-NTA purification, the N-terminal His_6_ tag was cleaved by the addition of 1.5 % Tobacco Etch Virus (TEV) protease, and the sample was dialyzed overnight into purification buffer C [20 mM Tris (pH 8), 150 mM NaCl, and 5 mM MgCl_2_]. The sample was then purified using the method previously described for the Mono Q anion exchange column, followed by further purification by gel filtration chromatography (HiPrep 26/60 Sephacryl S-200 HR; Amersham Biosciences). The collected sample was then dialyzed into purification buffer D [25 mM HEPES (pH 7.5), 50 mM KCl, 0.1 mM EDTA, 0.1 mM EGTA, and 2 mM DTT] overnight and concentrated using an Amicon Ultra-15 Centrifugal Filter Unit (Millipore) for storage.

The C-LiNK_TR_ construct was designed by replacing the 359-489 segment of C-LiNK with a stretch of six glycine residues. The corresponding cDNA, codon-optimized for *E. Coli* expression, was synthesized commercially (Genscript) and inserted into a pET-24a vector (containing a stop codon before the C-terminal His tag). The C-LiNK_TR_ protein was co-expressed with λ-phosphatase (pCDF-duet vector) in BL21(DE3) cells (New England Biolabs) and purified as previously described for eEF-2K_TR_ (18). The enzyme was incubated with λ-phosphatase (∼150 to 1 molar ratio) both during dialysis (overnight at 4 °C) and before injection in the gel filtration column (room temperature, 2 hours) in the presence of 1 mM MnCl_2_. The resulting enzyme lacked phosphorylation, as confirmed by ESI-QTOF mass spectrometry.

#### eEF-2

Eukaryotic elongation factor 2 (eEF-2) was purified from industrial yeast cake using a protocol similar to that of Jorgensen *et al.* (51). Yeast cells were suspended in purification buffer E [20 mM HEPES (pH 7.2), 10 % glycerol, 1 mM PMSF, and 1 mM DTT] containing 300 mM KCl and mechanically lysed using a bead beater. The lysate was centrifuged at 17,400 x g for 25 min. The supernatant was clarified by ultracentrifugation at 50,000 RPM (70 Ti fixed-angle rotor) for 60 min. The supernatant was dialyzed to remove KCl, then centrifuged at 17,400 x g for 20 min, and the resulting supernatant was filtered and applied to an SP FF 16/10 cation-exchange column. The sample was washed with 2 CV of purification buffer E containing 30 mM KCl and eluted over a 12 CV gradient up to 150 mM KCl. The eluted protein was applied to a Mono-Q column, washed with 2 CV of buffer E containing 40 mM KCl, and eluted over a 9 CV gradient up to 275 mM KCl. The collected fractions were re-applied to a Mono-Q column, washed with 2 CV of buffer E containing 40 mM KCl, and eluted over a 15 CV gradient up to 200 mM KCl. The protein was dialyzed into buffer F [25 mM HEPES (pH 7.5), 50 mM KCl, 2 mM DTT, and 5 mM MgCl_2_] overnight and concentrated using an Amicon Ultra-15 Centrifugal Filter Unit (Millipore) for storage.

### Multi-angle light scattering

Multi-angle light scattering experiments were performed as previously described (52). Experiments were performed at 25 °C on a DAWN HELEOS-II multiangle light scattering photometer attached to an Optilab T-rex refractive index detector and Wyatt QELS dynamic light scattering detector. Samples were loaded onto a TSK-GEL G300PWXL size exclusion column (7.8 mm x 300 mm with a pore size of 300 Å) through a Shimadzu LC-20AD HPLC system using buffer F as the solvent, with a flow rate of 0.4 mL/min. Molar mass (g/mol) was determined with 20 µL sample injections at a concentration of ∼20 µM. Astra 6 software (Wyatt Technology) was used to determine molar masses at the peak.

### Kinetic analyses

#### Manual mixing autophosphorylation assay

Enzyme (wild-type eEF-2K or C-LiNK) was incubated with CaM (CaM_WT_, CaM_N_, or CaM_C_) in assay buffer G [25 mM HEPES (pH 7.5), 50 mM KCl, 10 mM MgCl_2_, 100 µM EGTA, 150 µM CaCl_2_ (50 µM free Ca^2+^), 2 mM DTT, 20 µg/mL BSA, 0.005% Brij-35] with 0, 1, or 50 µM free Ca^2+^ at 30 °C for 10 min before initiating the reaction with 1 mM Mg^2+^•ATP. The reaction was quenched at specified time points by addition to 2.3 volumes of hot SDS-loading buffer [50 mM Tris (pH 6.8), 1.6 % SDS, 8% glycerol, 100 mM DTT, and 0.01 % bromophenol blue) and further incubated at 95 °C for 5 min. Samples were analyzed for autophosphorylation by western blotting using specific antibodies for eEF-2K (Santa Cruz Biotechnology) and its *p*T348, *p*S445, or *p*S500 forms (ECM Biosciences). To correct for sample loss and loading error, the phospho*-*antibody signal for each sample was divided by its corresponding total eEF-2K signal. Data were converted to fraction phosphorylated and fit to Eq. 1 for the apparent rate of autophosphorylation (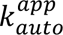).

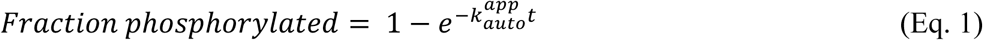

#### Rapid quench autophosphorylation assay

T348 autophosphorylation reactions were initiated by rapidly mixing equal volumes of solution A (400 nM eEF-2K with 4 µM CaM_WT_ or CaM_C_) and solution B (2 mM Mg^2+^•ATP) in assay buffer G with 150 µM CaCl_2_ (50 µM free Ca^2+^) at 30 °C on a KinTek RQF-4 apparatus. At specific time points, the reaction was quenched by the addition of buffer Q [200 mM KCl, 50 mM EDTA, and 10 mM EGTA in HEPES (pH 7.5)], then immediately dispensed into hot SDS-PAGE sample loading buffer and left at 95 °C for an additional 5 min. Samples were analyzed for incorporation of phosphate at T348 by western blotting. To correct for sample loss and loading error, the signal from the *p*T348 antibody for each sample was divided by its corresponding total eEF-2K signal. Data were converted to fraction phosphorylated and fit to Eq. 1 to obtain the rate of autophosphorylation (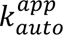).

#### General method for steady-state kinetic activity assays

*p*eEF-2K activity (WT or C-LiNK) was assayed at 30 °C in assay buffer G. Reactions were initiated by adding 1 mM [γ-^32^P]-ATP (100-1000 cpm/pmol) with 1 mM MgCl_2_. The activity was determined by calculating the rate of phosphate incorporation into a substrate peptide or yeast eEF-2.

Incorporation of phosphate into a peptide substrate (PepS, Ac-RKKYKFNED<u>T</u>ERRRFL-Amide) was measured by removing 10 µL aliquots and spotting onto P81 cellulose filter paper at 1 min intervals over a 5 min period. Spotted papers were immediately placed into 50 mM phosphoric acid. After completing the assay, all papers were washed 3 times in 50 mM phosphoric acid and once in acetone (10 min each wash), then dried. The amount of radioactivity associated with each paper was determined by measuring counts per minute (cpm) values on a Tri-Carb 2910 /TR liquid scintillation analyzer (Perkin Elmer).

Incorporation of phosphate into eEF-2 was measured by quenching reactions at 2 min with the addition of hot SDS-PAGE loading buffer followed by heating at 95 °C for an additional 5 min. Samples were then resolved by SDS-PAGE and stained using Coomassie Brilliant blue. The gels were dried, eEF-2 was excised, and the radioactivity was measured using a Tri-Carb 2910 /TR liquid scintillation analyzer (Perkin Elmer). The rate of phosphate incorporation (nmol/sec) was divided by enzyme concentration (nmol) to obtain *k*_obs_ (s^-1^) values.

#### CaM-dependence assays

Wild-type *p*eEF-2K dose-response assays were performed in assay buffer G using several CaM (CaM_WT_ or CaM_C_) concentrations in the presence or absence of 150 µM CaCl_2_. Assays performed without CaCl_2_ contained an additional 0.9 mM EGTA (1 mM total). Reactions against eEF-2 (5 µM) or peptide (150 µM) were performed with 0.5 or 1 nM *p*eEF-2K, respectively. Data were plotted as *k*_obs_ against the concentration of CaM and fit to Eq. 2 to obtain *k_obs,max_* (the maximal observed rate with saturating CaM under the specified conditions) and 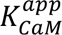 (the concentration of CaM required for half-maximal activity).

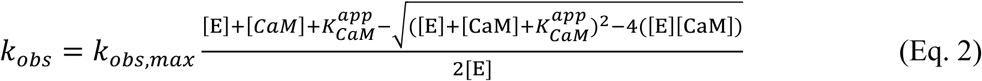

[E] is equal to the enzyme concentration in the assay, and [CaM] is the concentration of the CaM construct.

#### Substrate-dependence assays

The activity of 1 nM *p*eEF-2K (wild-type or C-LiNK) was measured in assay buffer G with 150 µM CaCl_2_ (50 µM free Ca^2+^) using several concentrations of PepS. Data were plotted as *k_obs_* against the concentration of PepS and fit to Eq. 3 to obtain best-fit values for 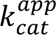 and 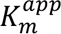.

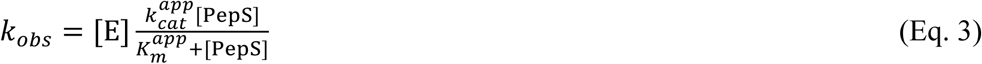

[E] is equal to the enzyme concentration in the assay, and [PepS] is the peptide substrate concentration.

#### Ca^2+^-dependence assays

The activity of 1 nM *p*eEF-2 (wild-type or C-LiNK) was measured in assay buffer G with an additional 0.9 mM EGTA (1 mM total) and different concentrations of CaCl_2_. The concentration of free CaCl_2_ in the assay was calculated using the Ca-Mg-ATP-EGTA calculator v1.0 (53), and data were plotted as *k_obs_* for the calculated concentration of free CaCl_2_ ([Ca^2+^]_free_).

### Mass spectrometric methods

Bottom-up digestion with LCMS/MS analysis was undertaken to identify the phosphorylation sites on C-LiNK. C-LiNK (50 µg) in 50 mM ammonium bicarbonate buffer (pH ∼8) was reduced with 5 mM DTT at 55 °C for 30 minutes, followed by alkylation with 15 mM iodoacetamide for 30 min in the dark before 5 mM DTT was added to stop alkylation. Trypsin was introduced at a 1:40 (w/w) protease:protein ratio, and the solution was incubated for 24 hours at 37 °C. The resulting proteolyzed solution was purified with a C18 micro spin column and eluted using 70% acetonitrile. The samples were dried using a SpeedVac and reconstituted in a solution containing 2% acetonitrile with 0.1% formic acid for subsequent LC-MS/MS analysis.

The digest was analyzed using a Dionex RSLC 3000 nano-LC system (Thermo Fisher Scientific) interfaced to an Orbitrap mass spectrometer. Approximately 200 ng (1 μL injection volume) of the digested protein was injected into an in-house packed 3 cm C-18 trapping column (3 μm, 300 Å pore size, 75 μm ID). Peptides were eluted into an in-house packed 20 cm C-18 analytical column (1.8 μm, 300 Å pore size, 75 μm, New Objective). Separation was performed using mobile phase A consisting of water and mobile phase B consisting of acetonitrile (both containing 0.1% formic acid). Peptide separation was achieved over a 60 min linear gradient of mobile phase B (2 to 90%) at a flow rate of 300 nL/min. MS analysis was performed using an Eclipse Orbitrap mass spectrometer (Thermo Scientific Instruments), and sample analysis was conducted in triplicate. MS1 spectra were collected with a resolution of 30,000 at 200 *m/z*. MS/MS spectra were acquired using the same resolution with a 25% normalized collision energy applied for collisional activation. Bottom-up data processing and analyses were performed with Byonic software (Protein Metrics, Cupertino, CA, USA). A mass tolerance limit of 10 ppm was used to validate both precursor and fragments. Posterior Error Probability scores and delta scores were used to evaluate the confidence of the identified peptides.

### Crystallization of C-LiNK_TR_ and structure determination

Crystals of C-LiNK were obtained using a 9.4 mg/mL protein solution in 20 mM Tris (pH 7.5), 100 mM NaCl, 1 mM TCEP, 1 mM ADP, and 0.35 mM CaCl_2_. The diffraction data were collected at the NSLS-II 17-ID-1 beamline on a crystal obtained by vapor diffusion by mixing 2 mL of protein with 1 mL of a 18.8% PEG3350 (Hampton Research) and 314 mM magnesium acetate solution (pH not adjusted further) in a 24 well-plate at room temperature. The data was processed using autoproc (54) and the structural model was generated using the phaser and the refine modules within the Phenix suite (55) and Coot (56).

### Cell-based assays

#### C-LiNK plasmid titration

0.15×10^6^ MCF10A^eEF-2K-/-^ cells (Sigma-Aldrich), maintained as described previously (without the use of penicillin or streptomycin) (19), were plated per well in a 35 mm plate. The following day, Lipofectamine 3000 (Invitrogen) was used to transfect cells with 0.1µg of wild-type eEF-2K pcDNA3 or 0.1µg, 0.5 µg, or 2 µg of C-LiNK pcDNA3 plasmid per well. All samples were transfected with a total of 2 µg of plasmid, with the difference between samples made up by adding empty pcDNA3 plasmid. Twenty-four hours later, cells were washed twice with cold PBS and lysed in M-PER lysis buffer and Halt™ protease and phosphatase inhibitor cocktail (Thermo Scientific). Lysates were cleared via centrifugation at 10,000 x g for 10 minutes, and the lysate protein concentration was analyzed via Bradford assay (Bio-Rad). As described below, 30 µg of lysate was subjected to SDS-PAGE and western blotting.

#### Cellular treatments

MCF10A^eEF-2K-/-^ cells were transfected as described above with either 0.5 µg empty pcDNA3, 0.1 µg wild-type eEF-2K pcDNA3 and 0.4 µg empty pcDNA3, or 0.5 µg C-LiNK pcDNA3. Transfected cells were treated with full media (no treatment, NT) or starved the following day by replacing media with Dulbecco’s phosphate-buffered saline (DPBS; starved, STRV) for two hours. Cells were then washed and lysed, and protein concentrations were determined as described above. Samples were subjected to western blot analysis as described below. 30 µg of lysate was analyzed by western blotting. Relative eEF-2 phosphorylation on Thr-56 was determined by normalization of *p*eEF-2 to total eEF-2 for each sample. Phosphorylated eEF-2 from the lysate of starved cells expressing wild-type eEF-2K (WT-STRV) was normalized to 1, and *p*eEF-2 levels from all samples were quantified relative to WT-STRV for each experiment.

### Protein detection by western blotting

30 µg of protein from cell lysates, as determined by the via Bradford assay (Bio-Rad) or a specified amount of recombinantly purified protein was resolved by SDS-PAGE, then transferred to PVDF membranes at 4 °C for 16 hours at 30 V in transfer buffer [25 mM Tris, 192 mM glycine, and 20% v/v methanol]. Membranes were then probed for total eEF-2K (C-12, Santa Cruz Biotechnologies), *p*T348 eEF-2K (ECM Biosciences), *p*445 eEF-2K (ECM Biosciences), *p*S359 eEF-2K (Invitrogen), total eEF-2 (Santa Cruz Biotechnologies), *p*Thr-56 eEF-2 (Cell Signaling), or CaM (C-lobe epitope, Cell Signaling). Primary antibodies were detected by corresponding goat anti-rabbit or anti-mouse secondary antibodies (LI-COR).

Additional experimental procedures and protocols are available in the Supporting Information.

## Data availability

The structure factors and coordinates have been deposited in the Protein Data Bank (PDB) with accession code 9OCW. All other data are available upon request to the corresponding authors.

## Supporting information

This article contains supporting information.

## Acknowledgments

This work is supported by NIH award R01 GM123252 (to KND and RG), NIH (R35GM13965, JSB), the Robert A. Welch Foundation (F-1155, JSB), and a CPRIT award RP210088 (to KND). EYW was supported by the LEADER Program, College of Pharmacy, The University of Texas, Austin. Use of the NYX beamline (19-ID) at the National Synchrotron Light Source II (NSLS-II) is supported by the member institutions of the New York Structural Biology Center. NSLS-II is a United States Department of Energy (DOE) Office of Science user facility operated for the DOE Office of Science by Brookhaven National Laboratory under Contract DE-SC0012704.

## Author contributions

Kimberly J. Long: Conceptualization, Investigation, Writing-original draft, Writing-review & editing. Luke S. Browning: Investigation, Writing-review & editing. Andrea Piserchio: Investigation, Writing-review & editing. Eta A. Isiorho: Investigation. Mohamed Gadallah: Investigation, Writing-review & editing. Jomai Douangvilay: Investigation. Elizabeth Y. Wang: Investigation. Justin K. Kalugin: Writing-review & editing. Jennifer S. Brodbelt: Writing-review & editing. Ranajeet Ghose: Conceptualization, Writing-review & editing. Kevin N. Dalby: Conceptualization, Writing-review & editing

## Abbreviations

1-PP: lamda protein phosphatase
apo-CaM: Ca^2+^-free CaM
C-LiNK: CaMC linked to ΔN eEF-2K
C-LiNK: C-LiNK with the 359-489 segment of C-LiNK replaced by 6 glycines
CaM: calmodulin
CaMC: CaM truncation containing residues 76-148
CaMN: CaM truncation containing residues 1-80
CaMWT: wild-type CaM
CTM: CaM targeting motif
eEF-2K: eukaryotic elongation factor 2 kinase
eEF-2K_TR_: eEF-2K truncated construct containing residues 71-358 connected to residues 490-725 by a 6-glycine linker
eEF-2: eukaryotic elongation factor 2
ESI-QTOF: electrospray ionization quadrupole time of flight mass spectrometry
LC-MS/MS: liquid chromatography tandem mass spectrometry
MCF10A^eEF-2K-/-^: homozygous eEF-2K knock out in non-tumorigenic epithelial cell line
NMR: nuclear magnetic resonance
*p*eEF-2K: T348 phosphorylated eEF-2K
PepS: eEF-2K peptide substrate (Ac-RKKYKFNEDTERRRFL-Amide)
TEV: protease Tobacco Etch Virus protease

## Supporting information

**Fig. S1.**
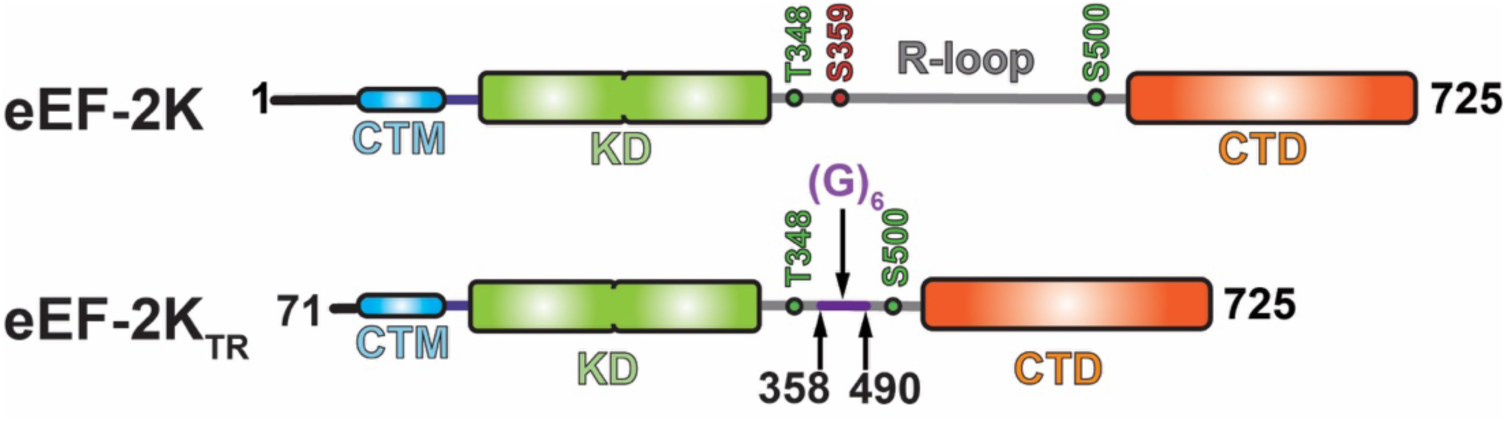
Domain structure of eEF-2K. Full-length eEF-2K consists of 725 residues organized in an N-terminal calmodulin targeting motif (CTM), an α−kinase domain (KD), a regulatory loop (R-loop) containing multiple phosphorylation sites that terminates in an α−helical C-terminal domain (CTD). The stimulatory autophosphorylation sites, T348 and S500, and the inhibitory phosphorylation site, S359, discussed in the text, are indicated in green and red, respectively. The truncated eEF-2K construct (eEF-2K_TR_) is missing 70 N-terminal residues, and 6 glycines have replaced the segment of the R-loop between 359 and 489.

**Fig. S2.**
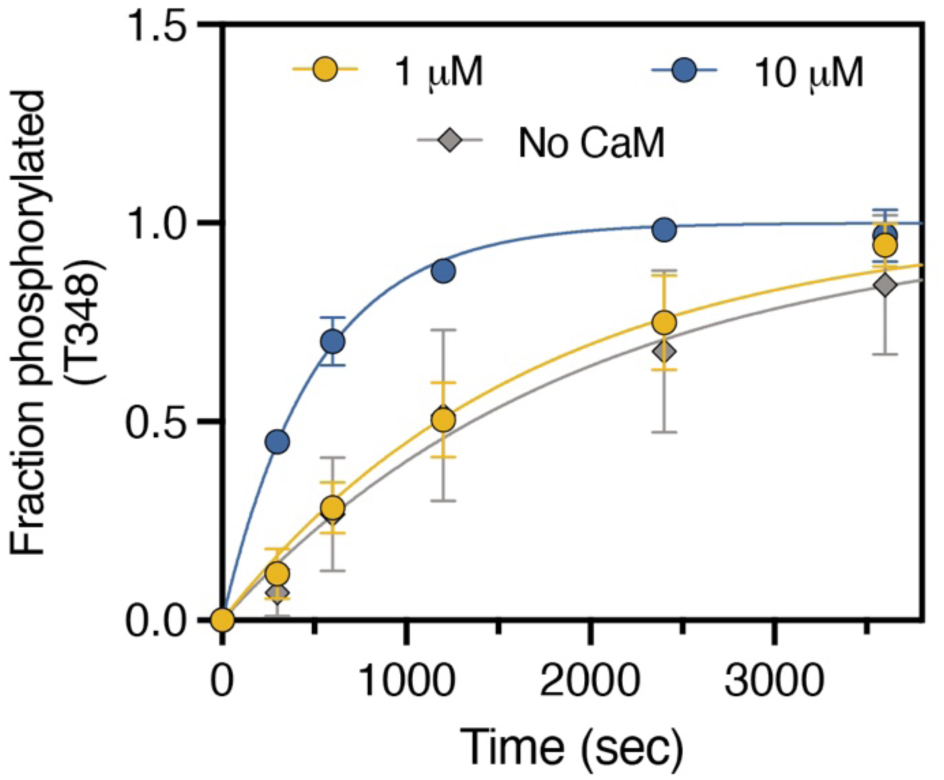
Measurement of the ability of isolated CaM_N_ to drive auto-phosphorylation on T348. CaM_N_ and no CaM samples (measured in duplicate) from Fig. 2A were quantified and plotted as the fraction of T348 phosphorylation over time (min) and fit to Eq. 1. The best-fit 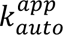 values for 1 µM CaM_N_ = 0.0006 ± 0.00004 s^-1^; 10 µM CaM_N_ = 0.002 ± 0.00009 s^-1^; no CaM= 0.0005 ± 0.00007 s^-1^.

**Fig. S3.**
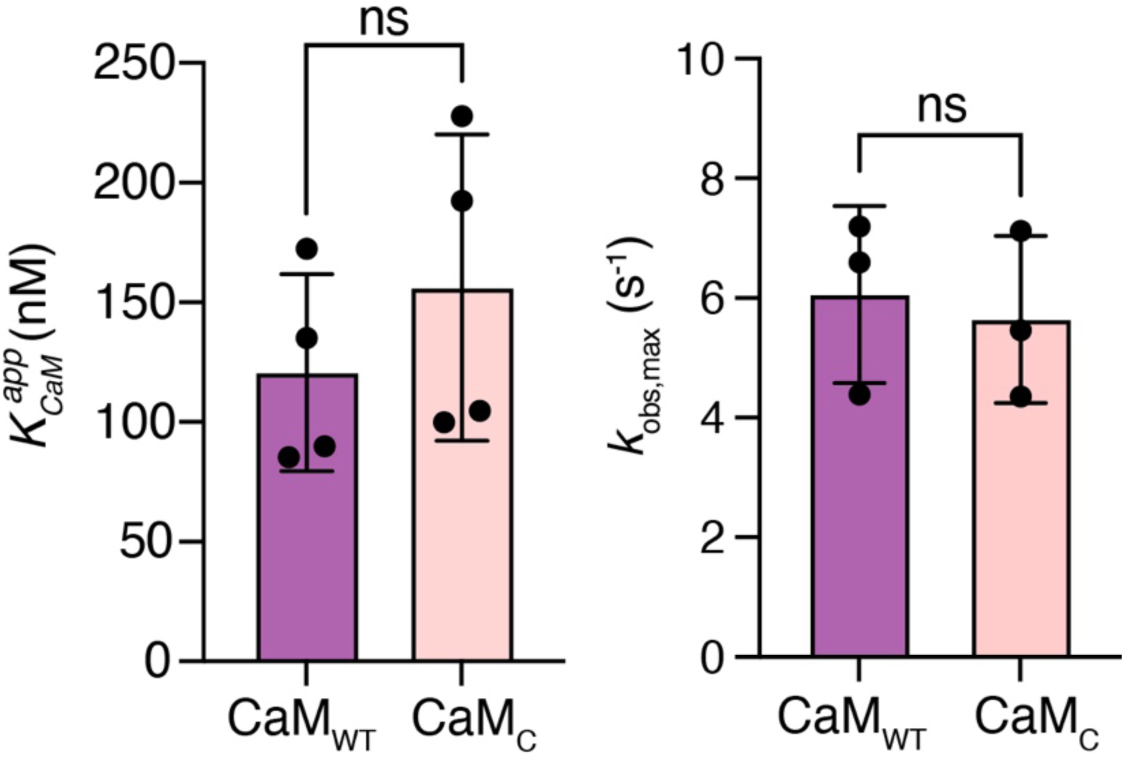
Quantification of *p*eEF-2K CaM dependence against yeast eEF-2. Samples (depicted in Fig. 3B) were extracted from the gel and quantified by scintillation counting. The *k*_obs_ was plotted against the CaM concentration, and data were fitted to (Eq. 2) to obtain best-fit values for *k*_obs,max_ and 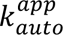. The average *k*_obs,max_ (CaM_WT_ = 6.1 ± 1.5 s^-1^; CaM_C_ = 5.6 ± 1.4 s^-1^) and 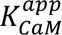 (CaM ± 41 s^-1^; CaM_C_ = 156 ± 64 s^-1^) values were similar _WT_ = 121 _C_ in the two cases. Individual values obtained from independent measurements are represented by the filled circles; the average and standard deviation over the measurements are depicted by the filled bars and vertical lines, respectively. An unpaired *t*-test indicated no significant (ns) difference in the *k*_obs_,_max_ (P= 0.74) or 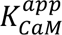 (P= 0.39) values between the two cases.

**Fig. S4.**
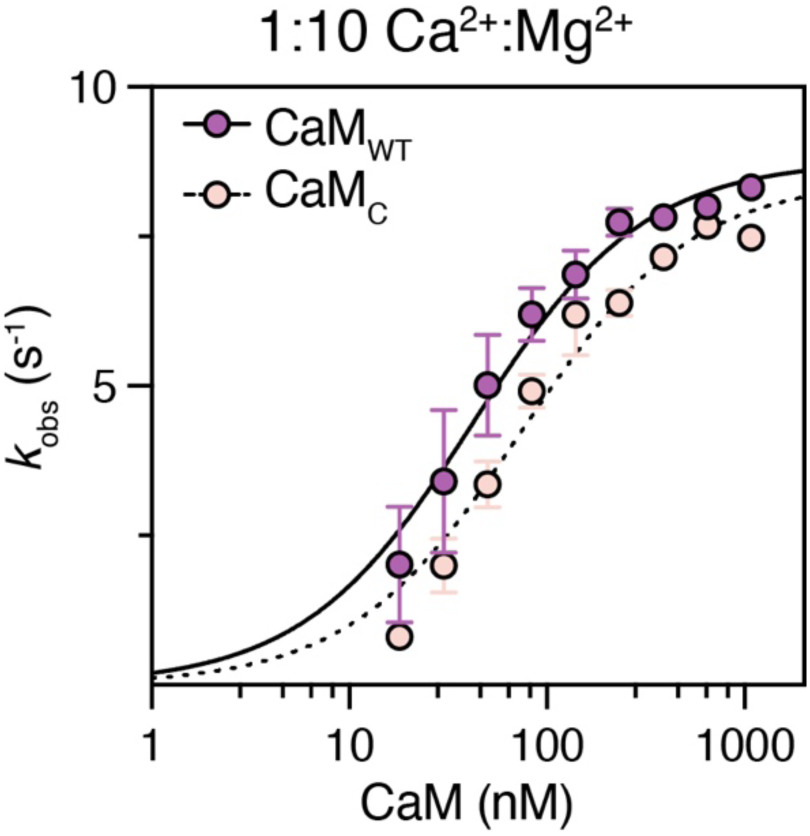
*p*eEF-2K activity dependence on CaM_WT_ or CaM_C_ at a Ca^2+^ : Mg^2+^ ratio of 1:10. The dose-dependence of *p*eEF-2K (1 nM) activity on CaM_WT_ or CaM_C_ was measured against 150 µM PepS with 1 mM [γ-^32^P]-ATP in the presence of 1 mM free Ca^2+^ and 10 mM free Mg^2+^. The *k*_obs_ values were plotted against the concentration of CaM, and data were fit to Eq. 2 to obtain best-fit values for *k*_obs,max_ (CaM_WT_ = 8.8 ± 0.3 s^-1^; CaM_C_ = 8.5 ± 0.3 s^-1^) and 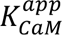 (CaM_WT_ = 41 ± 5 nM; CaM_C_ = 73 ± 9 nM). Compare 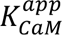 values for’ CaM_WT_ (67 ± 7 nM) and CaM_C_ (85 ± 10 nM) at a Ca^2+^ : Mg^2+^ ratio of 1:200.

**Fig. S5.**
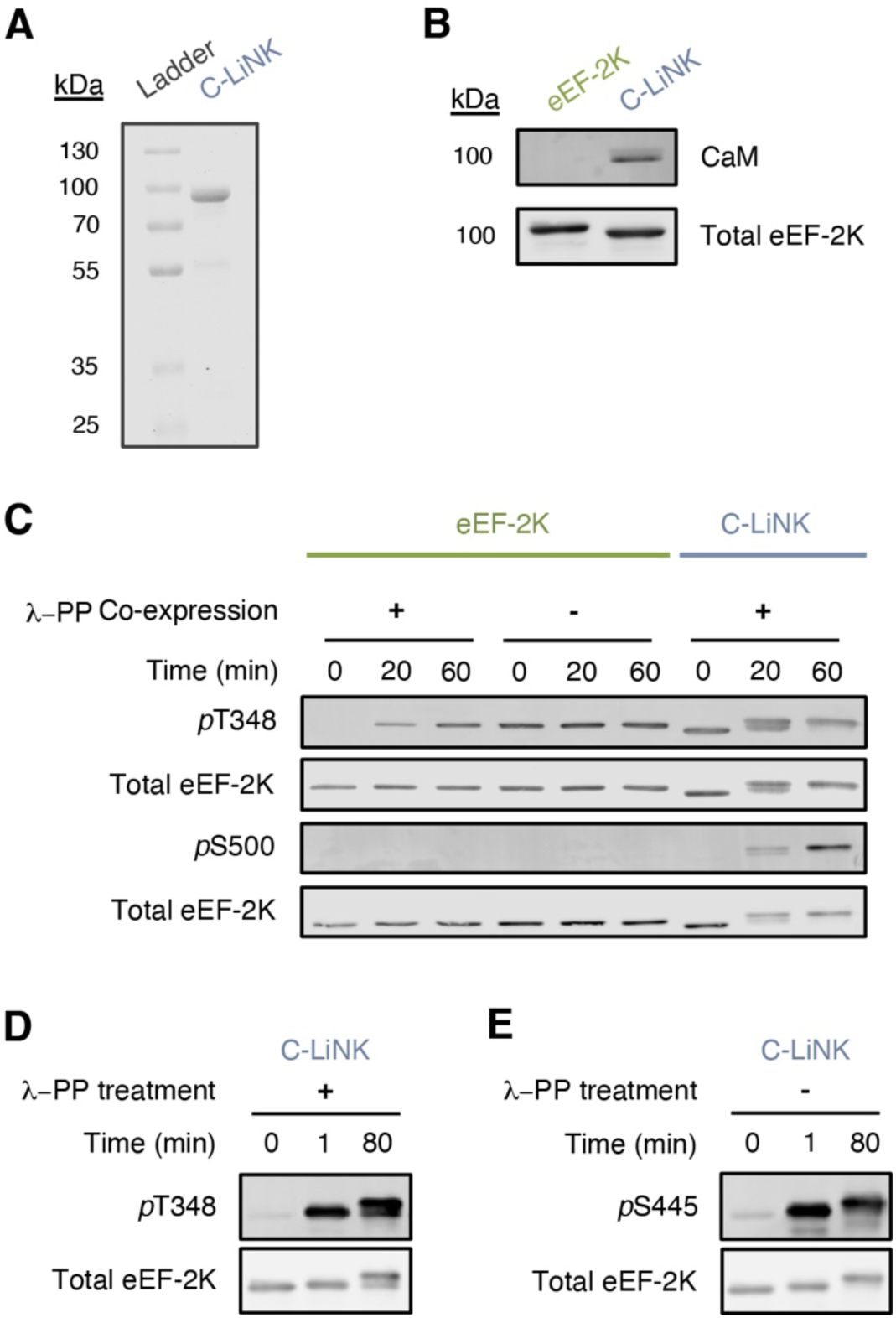
Autophosphorylation of C-LiNK. (**A**) 2 µg of purified C-LiNK protein was run on a 10% polyacrylamide denaturing gel and stained for total protein content with Coomassie blue. (**B**) 0.5 µg of wild-type eEF-2K or C-LiNK protein were probed using eEF-2K and CaM-specific antibodies that recognize the C-termini of eEF-2K (Santa Cruz, residues 436-725) and CaM (Cell Signaling) (**C**) Purified wild-type eEF-2K (+/-co-expression with λ-phosphatase) or C-LiNK (co-expressed with λ-phosphatase) were incubated with 1 mM ATP for 0, 30, or 60 mins in the presence of 50 µM Ca^2+^ before quenching with hot SDS-loading buffer. Samples were subject to western blotting to detect total enzyme and phosphate incorporation through autophosphorylation at T348 and S500 (0.125 µg of protein was loaded). (**D-E)** C-LiNK was expressed in the presence of λ-phosphatase and either treated or not during the purification process with λ-phosphatase (+/-λ-PP treatment). The autophosphorylation reaction was performed in assay buffer H [25 mM HEPES (pH 7.5), 50 mM KCl, 10 mM MgCl_2_, 100 µM EGTA, 2 mM DTT, 20 µg/mL BSA, 0.005% Brij-35] at 30 °C with 200 nM enzyme in the presence of 50 µM free CaCl_2_. The reaction was initiated with 1 mM ATP, then quenched at various time points by adding hot SDS loading buffer. 150 ng of enzyme was run on SDS-PAGE, and phosphate incorporation was detected by western blotting for (**D**) *p*T348 or (**E**) *p*S445 and (**D-E**) total eEF-2K.

**Fig. S6.**
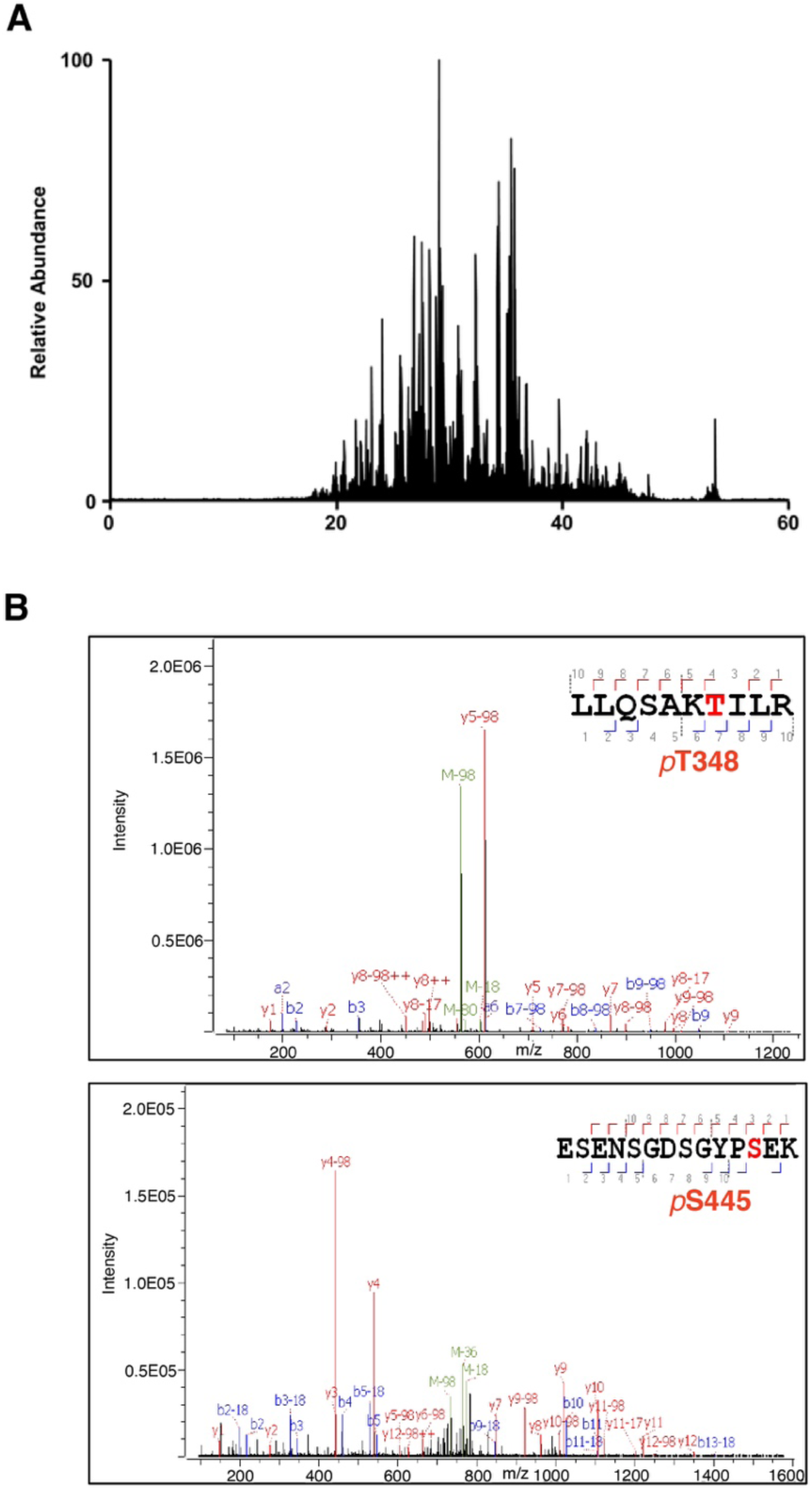
Expressed C-LiNK is phosphorylated on T348. **(A)** Total ion chromatogram showing peptide separation throughout the LC-MS/MS run. **(B)** CID fragmentation pattern of one peptide ^342^LLQSAK*p***T**ILR^351^ (2+, upper spectrum), indicative of phosphorylation on T348. Also shown (lower spectrum) is the fragmentation pattern of a second peptide ^434^ESENSGDSGYP*p***S**EK^447^ (2+), indicating the presence of phosphorylation on S445. The former is highly abundant (> 99%), while the latter has low abundance (1%). The complete lists of assigned fragment ions are summarized in Tables S1 and S2.

**Fig. S7.**
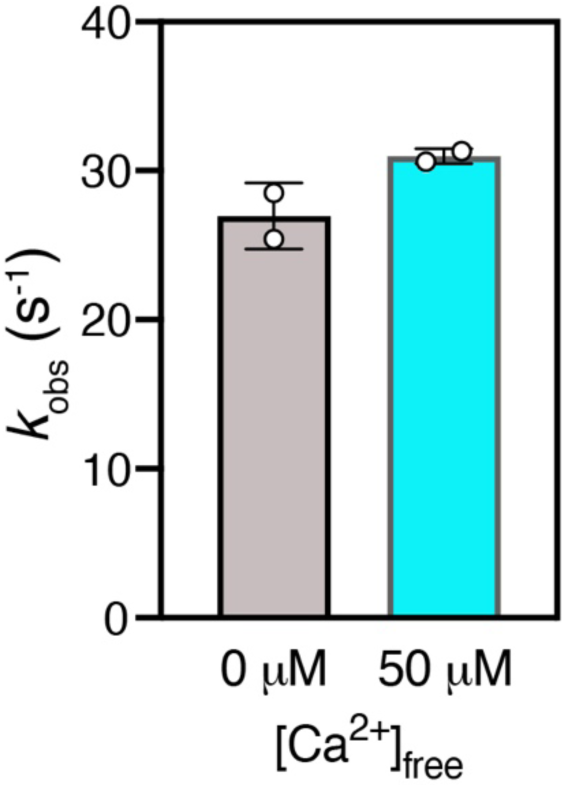
Influence of Ca^2+^ on the activity of C-LiNK. 23 µM C-LiNK in storage buffer [25 mM HEPES (pH 7.5), 50 mM KCl, 0.1 mM EGTA, 0.1 mM EDTA, 2 mM DTT, and 10% glycerol] was dialyzed for 19 hours against assay buffer H [25 mM HEPES (pH 7.5), 50 mM KCl, 10 mM MgCl_2_, 100 µM EGTA, 2 mM DTT, 20 µg/mL BSA, 0.005% Brij-35] with 0.9 mM EGTA added. Next, the activity of 4 nM C-LiNK in assay buffer H containing 0.9 mM EGTA was measured against 150 µM PepS. CaCl_2_ was added to the indicated reaction to a concentration of 50 µM free Ca^2+^, and MgCl_2_ was added to each reaction to result in 10 mM free Mg^2+^. Reactions were initiated by adding 1 mM [γ-^32^P]-ATP.

**Fig. S8.**
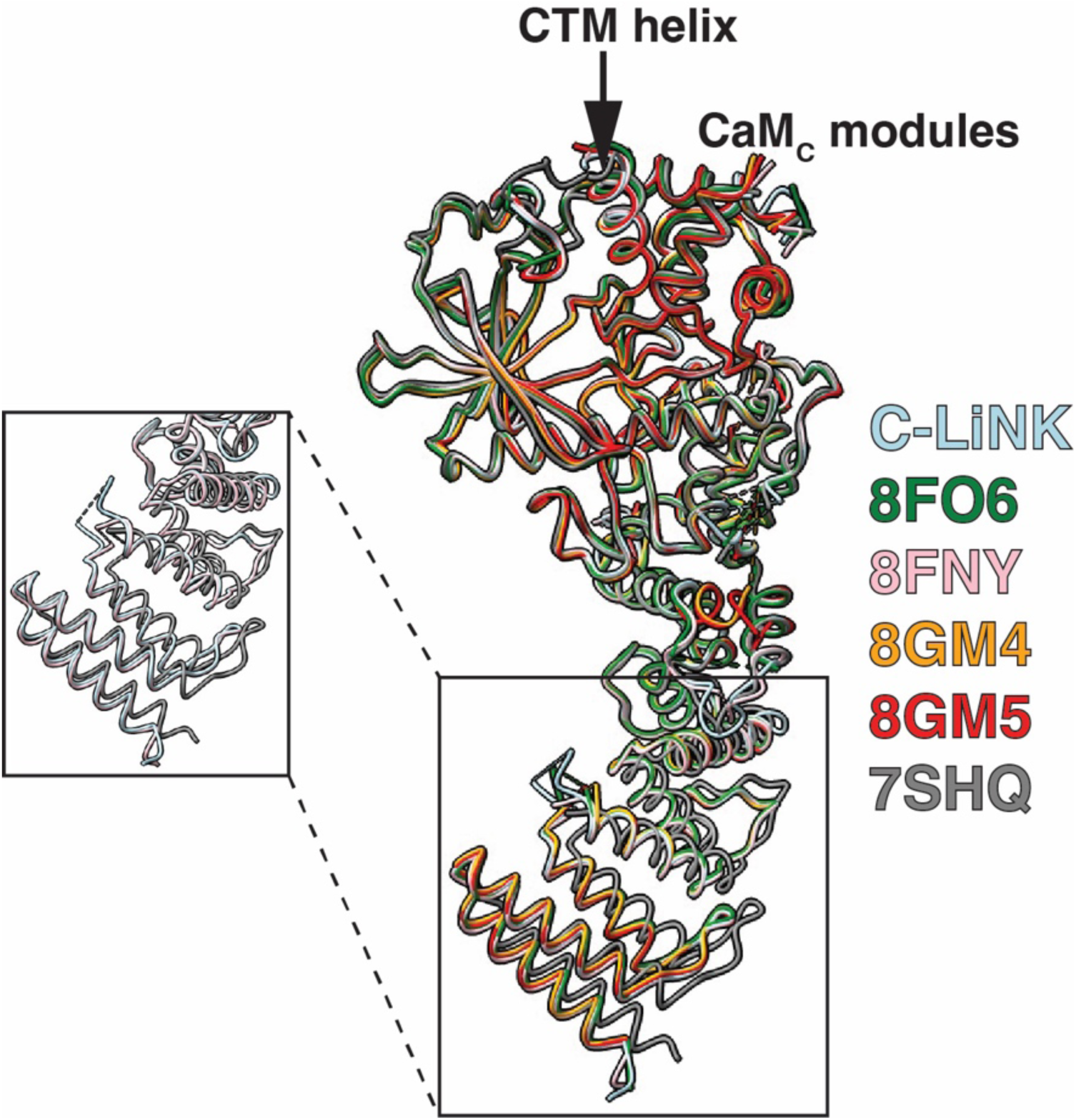
C-LiNK_TR_ has a similar overall conformation as the structural modules of the CaM•*p*eEF-2K_TR_ complex. An overlay of all available CaM•*p*eEF-2K_TR_ complex structures with C-LiNK_TR_ suggests that the overall conformations of its eEF-2K_TR_ and the CaM_C_ modules remain unchanged. The 7SHQ structure shows a slightly different CTD conformation than all other structures, including C-LiNK_TR_ (see inset for an expanded view; the CTDs of C-LiNK_TR_, 8FNY, and 7SHQ are shown). The 7SHQ structure was solved using different crystallization conditions, and its slightly altered CTD conformation is likely a reflection of interdomain flexibility.

**Fig. S9.**
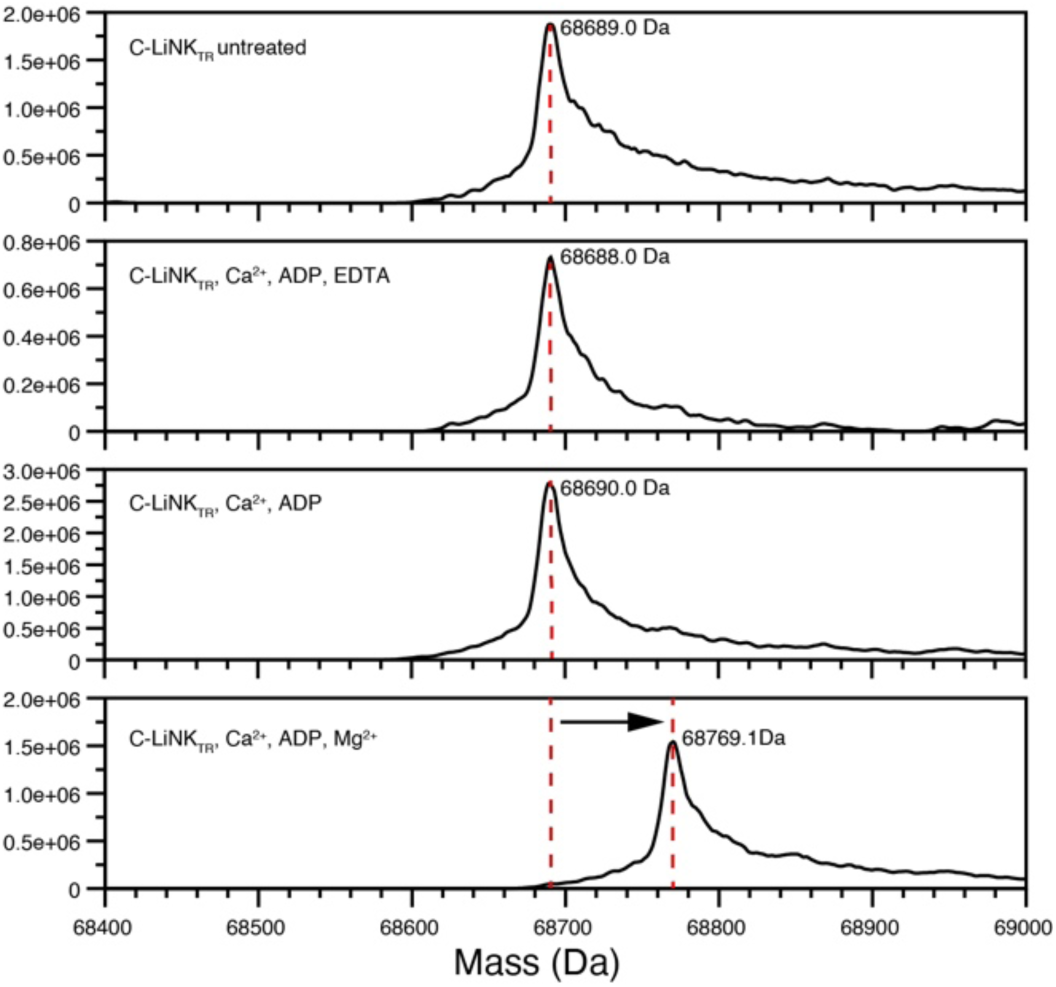
Intact mass measurements obtained for expressed C-LiNK_TR_ using an ESI-Q-TOF mass spectrometer. (Top panel) C-LiNK_TR_ co-expressed with λ-PP and further incubated with λ-PP as described in the main text. (Second panel from the top) Protein stock (9.4 mg/mL) used for crystallography in a buffer containing 20 mM Tris (pH 7.5), 100 mM NaCl, 1 mM TCEP, 1 mM ADP, and 0.35 mM CaCl_2_ incubated with 20 mM EDTA. (Third panel from the top) Protein stock solution used for crystallography without EDTA incubation. (Bottom panel) Protein stock used for crystallography incubated with 10 mM MgCl_2_ for 24 hours at 4 °C. The last shows a shift in mass equivalent to the addition of a single phosphate upon incubation with Mg^2+^.

**Fig. S10.**
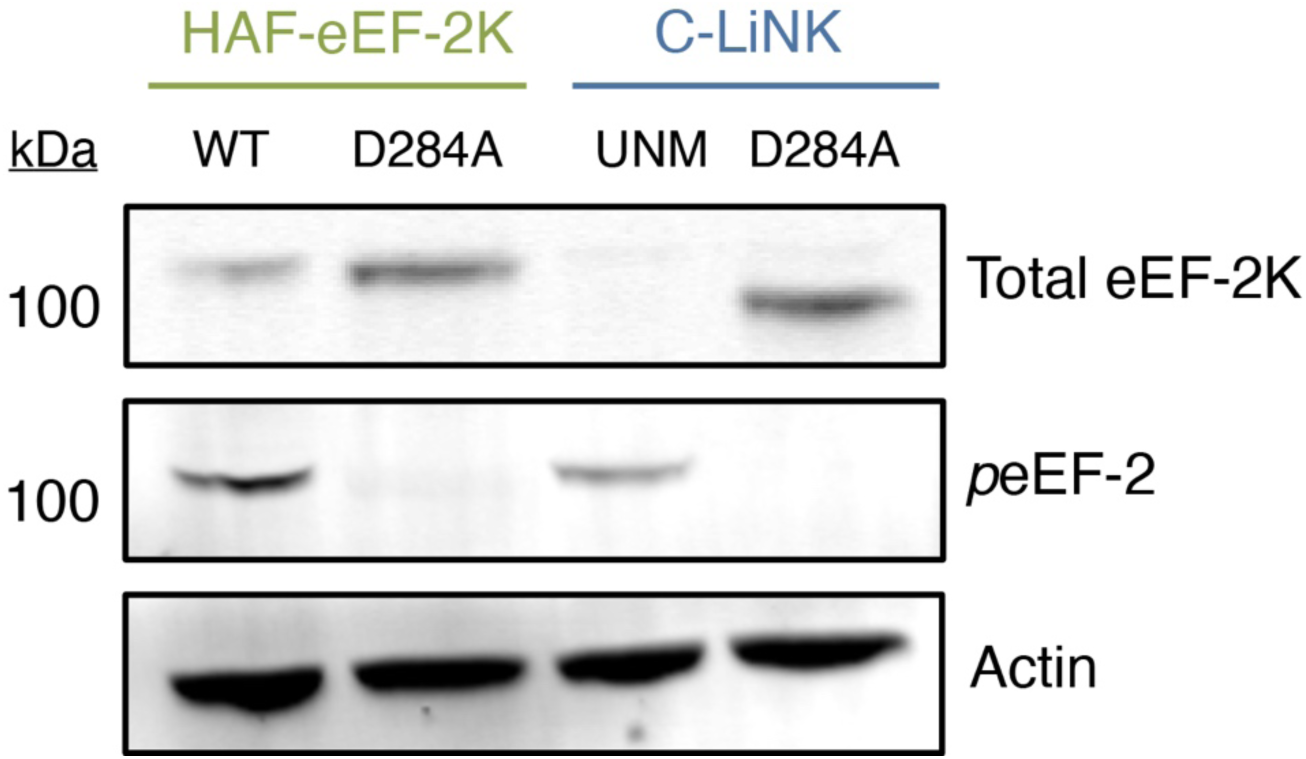
Mutation of a C-LiNK catalytic residue rescues protein levels. MCF10A^eEF-2K-/-^ cells were transfected with 0.1 µg of either wild-type (WT) or D284A HAF-eEF-2K, unmodified (UNM, D284) or D284A C-LiNK. After 16 hours, cells were lysed and 40 µg of cell lysate was analyzed by western blotting using specific antibodies for eEF-2K, *p*eEF-2, and actin (loading control).

**Fig. S11.**
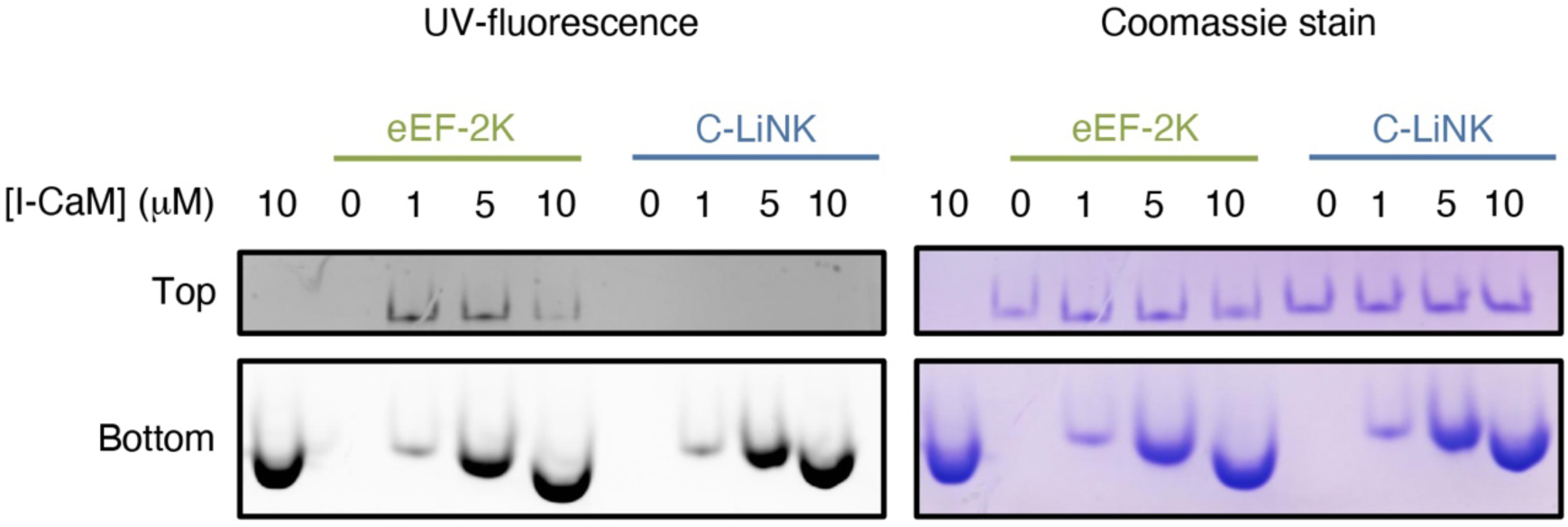
Exogenous CaM does not bind C-LiNK. 400 nM eEF-2K was incubated with the indicated concentrations of CaM labeled with the fluorescent dye IAEDANS (I-CaM; the dye was covalently attached to position 75 in a K75C mutant of CaM) and 100 µM free CaCl_2_ for 20 min at room temperature before running samples on a gradient (4-15%) native gel. Fluorescence was detected by exposing the gel to ultraviolet light for 0.5 (bottom) or 34.5 (top) seconds before capturing the image. The gel was Coomassie stained to detect total protein.

## Additional experimental procedures

### Effects of increased Ca^2+^ to Mg^2+^ ratio on eEF-2K activity

1 nM *p*eEF-2K activity was measured in buffer H (see main text) with 1.15 mM CaCl_2_ (1 mM free Ca^2+^) with varied CaM_WT_ or CaM_C_. Data were fit to Eq. 1.

### Dephosphorylation of C-LiNK

To further dephosphorylate C-LiNK protein co-expressed with a plasmid containing λ-PP, it was treated with λ-PP during purification. The cleared cell lysate was incubated with Ni-NTA beads (Qiagen) for 1 hour at 4 °C. In a chromatography column, the beads were washed with purification buffer containing 25 mM HEPES (pH 8), 100 mM NaCl, 20 mM Imidazole, and 0.03% Brij-30 (v/v), 1% 2-mercaptoethanol (v/v), 0.1 mM TPCK, 0.1 mM PMSF, and 1 mM benzamidine. The tagged C-LiNK was then eluted in 30 mL of a buffer containing 25 mM HEPES (pH 8), 250 mM Imidazole, and 0.03% Brij-30 (v/v), 1% 2-mercaptoethanol (v/v), 0.1 mM TPCK, 0.1 mM PMSF, and 1 mM benzamidine. λ-PP (New England Biolabs) was added to the eluted protein at a 1:250 ratio with 1 mM MnCl_2_. The sample was dialyzed against purification buffer with 1 mM MnCl_2_ for 16 hours at 4 °C. Following dialysis, the sample was re-purified through the Ni-NTA column before treating with TEV protease and continuing successive chromatographic steps.

### Autophosphorylation of C-LiNK

C-LiNK was expressed in the presence of λ-PP. Further treatment of the enzyme with λ -PP is discussed above. The autophosphorylation reaction was performed in assay buffer H at 30 °C with 200 nM enzyme in the presence of 50 µM free CaCl_2_. The reaction was initiated with 1 mM ATP, then quenched at various time points by adding hot SDS loading buffer. 150 ng of phosphorylated enzyme was run on SDS-PAGE, and phosphate incorporation was detected by western blotting for *p*T348 (ECM Biosciences) or *p*S445 (ECM Biosciences).

### Effect of extended Ca^2+^ chelation on C-LiNK activity

Approximately 23 µM C-LiNK in storage buffer [25 mM HEPES (pH 7.5), 50 mM KCl, 0.1 mM EGTA, 0.1 mM EDTA, 2 mM DTT, and 10% glycerol] was dialyzed for 19 hours against dialysis buffer containing 25 mM HEPES (pH 7.5), 50 mM KCl, 10 mM MgCl_2_, 1 mM EGTA, 2 mM DTT, 0.005% Brij-35. Subsequently, 4 nM dialyzed C-LiNK activity was measured against 150 µM PepS in dialysis, with or without CaCl_2_ (50 µM free Ca^2+^), and MgCl_2_ (10 mM free Mg^2+^). Reactions were initiated with 1 mM [γ-^32^P], and time points were taken over 5 min.

### Native protein detection

400 nM eEF-2K was incubated with indicated concentrations of IAEDANS-labeled CaM_K75C_ (I-CaM) in native buffer containing 5 mM HEPES (pH 6.8), 50 mM KCl, 100 µM EGTA, 150 µM CaCl_2_, and 0.005% Brij for 20 min at room temperature before running samples on a gradient (4-15%) Mini-Protean TGX native gel (BioRad) using Tris/glycine native running buffer (pH 8.3; BioRad).

**Table S1.** Fragment ion assignments for phosphorylated peptide LLQSAK**T**ILR (*m/z* 611.85, 2+, retention time: 26.44 min) identified by LC-MS/MS-HCD following tryptic digestion of C-LiNK. Theoretical and observed masses of b- and y-ions are listed along with the corresponding mass differences and errors (ppm)

| **Ion Type** | **Number of residues** | **Theoretical Mass (Da)** | **Observed Mass (Da)** | **Mass difference (Da)** | **Mass Error (ppm)** |
| --- | --- | --- | --- | --- | --- |
| a | 2 | 199.1805 | 199.1799 | -0.0006 | -3.0 |
| b | 2 | 227.1754 | 227.1749 | -0.0005 | -2.2 |
| b | 9 | 1048.5802 | 1048.5698 | -0.0104 | -9.9 |
| y | 1 | 175.119 | 175.1186 | -0.0004 | -2.3 |
| y | 6 | 781.4332 | 781.4294 | -0.0038 | -4.9 |
| y | 7 | 868.4652 | 868.4613 | -0.0039 | -4.5 |
| y | 8 | 996.5238 | 996.5217 | -0.0021 | -2.1 |
| y (2+) | 8 | 498.7655 | 498.7642 | -0.0013 | -2.6 |
| y (2+) | 9 | 555.3075 | 555.306 | -0.0015 | -2.7 |

Number of residues indicates the number of amino acids contained in the fragment ion. For a- and b-type ions, this number is anchored relative to the N-terminus. For y-type ions, this number is anchored relative to the C-terminus. All ions are singly charged except for two doubly charged y-type ions.

**Table S2.**
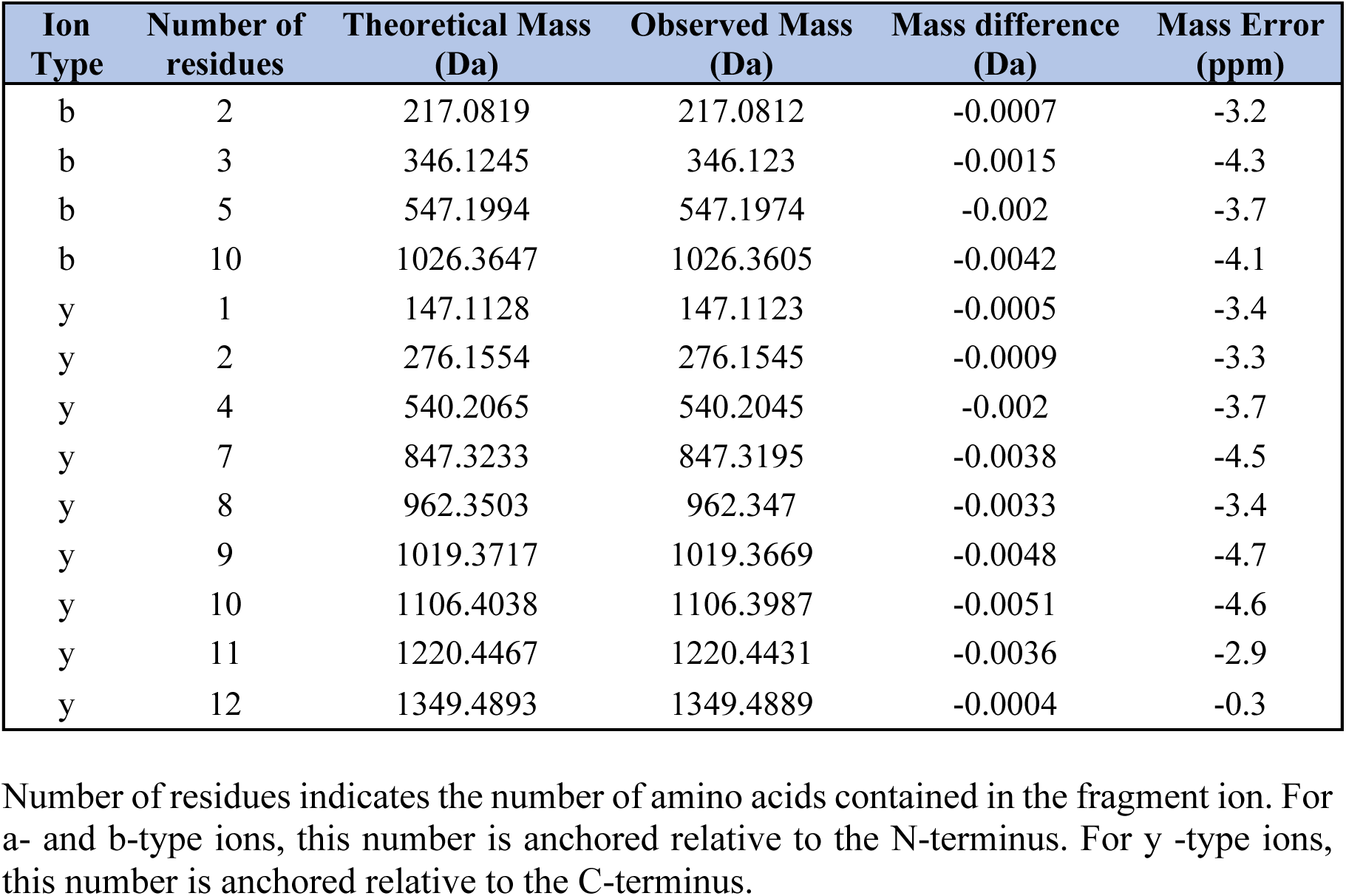
Fragment ion assignments for phosphorylated peptide ESENSGDSGYPSEK (*m/z* 783.28, 2+, retention time: 18.54 min) identified by LC-MS/MS-HCD following tryptic digestion of C-LiNK. Theoretical and observed masses of b-and y-ions are listed along with the corresponding mass differences and errors (ppm).

| **Ion Type** | **Number of residues** | **Theoretical Mass (Da)** | **Observed Mass (Da)** | **Mass difference (Da)** | **Mass Error (ppm)** |
| --- | --- | --- | --- | --- | --- |
| b | 2 | 217.0819 | 217.0812 | -0.0007 | -3.2 |
| b | 3 | 346.1245 | 346.123 | -0.0015 | -4.3 |
| b | 5 | 547.1994 | 547.1974 | -0.002 | -3.7 |
| b | 10 | 1026.3647 | 1026.3605 | -0.0042 | -4.1 |
| y | 1 | 147.1128 | 147.1123 | -0.0005 | -3.4 |
| y | 2 | 276.1554 | 276.1545 | -0.0009 | -3.3 |
| y | 4 | 540.2065 | 540.2045 | -0.002 | -3.7 |
| y | 7 | 847.3233 | 847.3195 | -0.0038 | -4.5 |
| y | 8 | 962.3503 | 962.347 | -0.0033 | -3.4 |
| y | 9 | 1019.3717 | 1019.3669 | -0.0048 | -4.7 |
| y | 10 | 1106.4038 | 1106.3987 | -0.0051 | -4.6 |
| y | 11 | 1220.4467 | 1220.4431 | -0.0036 | -2.9 |
| y | 12 | 1349.4893 | 1349.4889 | -0.0004 | -0.3 |

Number of residues indicates the number of amino acids contained in the fragment ion. For a- and b-type ions, this number is anchored relative to the N-terminus. For y -type ions, this number is anchored relative to the C-terminus.

